# Pop-Corn: Predicting Perturbation Phenotype Effects Across Single-Cell and Spatial Contexts

**DOI:** 10.64898/2026.09.25.754560

**Authors:** Jiapeng Chen, Yan Cui, Yanjun Shao, Na Sun, María Rodríguez Martínez

**Author notes:** Contributing authors. These authors contributed equally to this work (Jiapeng Chen, Yan Cui).

## Abstract

Genetic perturbations can reshape cell populations by altering the relative abundance of specific cell types and states within the profiled population, including increases, decreases and states that become detectable after perturbation. Pooled single-cell screens, such as Perturb-seq, measure such responses at scale. However, only a small fraction of possible perturbations can be tested experimentally. A central challenge is therefore to predict compositional shifts induced by unseen perturbations. Many perturbation-prediction methods do not directly optimize for this outcome; instead, they predict gene-expression responses and infer cell-type and cell-state composition downstream. Surprisingly, we find that even models that accurately predict perturbation-induced changes in average gene expression perform poorly at forecasting these compositional shifts. To address this gap, we present Pop-Corn, a method that directly predicts how a perturbation reshapes cell-type composition without reconstructing gene expression. In the primary T-cell benchmark, Pop-Corn predicted the overall cell-state composition of held-out perturbations more accurately than the evaluated expression-prediction pipelines, while better preserving the diversity of observed cell states. We further extend Pop-Corn to intact tissue, where it predicts perturbation-induced cell-type proportion changes in local cellular neighborhoods and uses attention patterns to generate hypotheses about context-dependent cellular interactions. Retrospective virtual screens support the use of Pop-Corn to prioritize perturbations for experimental follow-up according to their predicted effects on cell-state composition.

## Introduction

Perturb-seq measures the transcriptional consequences of genetic perturbations at single-cell resolution, revealing increases and decreases in the relative abundance of cell types and states within the profiled population, as well as states that become detectable after perturbation [1, 2]. We refer to these changes in population composition as perturbation phenotypes, distinguishing them from transcriptional responses within individual cells. Yet even pooled screens sample only a small fraction of the genome-wide singlegene and combinatorial intervention space. Predictive models are therefore needed to prioritize unmeasured perturbations for experimental follow-up [3–9], therapeutic discovery and mechanistic investigation [10, 11]. Predicting these population changes poses three challenges. First, sparse and noisy single-cell measurements complicate cell-state annotation and its transfer across perturbation conditions. Composition prediction therefore depends on the supplied annotations; Pop-Corn uses these labels rather than learning a new annotation system. Second, models must capture heterogeneous responses across the population. Third, prediction for unseen perturbations requires biological information that relates new targets to those observed during training.

Most existing methods, however, formulate and evaluate this inherently population-level problem as the reconstruction of post-perturbation gene expression profiles [3–5, 8, 12–14]. Recent benchmarks question whether strong transcriptional prediction scores necessarily reflect accurate perturbation-specific responses [14–16]. Predictions close to an average perturbed profile or to control can obscure differences between perturbations. Recovering these differences and preserving the distribution of cell states are distinct requirements for phenotype prediction. Errors in either may limit phenotype-based prioritization and experimental design.

Population composition is itself biologically consequential: perturbation-induced changes in cell identity, state abundance and rare-state emergence reveal gene function, resolve genetic interactions and identify regulators of therapeutically relevant cellular programs [2, 10, 17, 18]. The relevant prediction target is therefore not transcript-level fidelity alone, but the distribution of cell types and states produced by a perturbation. Recent methods have begun to shift towards phenotypic prediction. CellFlow models perturbation-conditioned single-cell distributions, but still derives population composition indirectly from generated cell states [5]. Prophet predicts aggregate experimental phenotypes, such as drug sensitivity and whole-embryo cell-type frequencies, without resolving the single-cell context underlying these readouts [6]. A general framework that directly links perturbations to population-level phenotypic change is therefore still lacking.

Here we reformulate perturbation prediction as the direct prediction of cell-type composition. Pop-Corn aggregates control single-cell states with perturbation representations to predict population composition under unseen perturbations, without reconstructing gene expression. This formulation addresses dissociated screens, in which tissue architecture is lost. In intact tissues, however, perturbation phenotypes may also depend on the local cellular context. We therefore extend Pop-Corn to spatial perturbation experiments [19], enabling context-aware prediction of niche-level composition and using attention to generate hypotheses about neighbourhood-associated perturbation effects [20]. Across these settings, Pop-Corn provides a direct route from control-cell profiles and perturbation identity to predicted cell-state proportions. By predicting changes in the relative abundance of annotated cell states, these outputs directly guide biologists in selecting perturbations and cell-state readouts for experimental follow-up.

## Results

### Expression-level prediction fails to capture phenotypic shifts

To determine whether existing perturbation prediction methods recover biologically meaningful phenotypic outcomes, we benchmarked four state-of-the-art expression-centric models using the primary human T cell Perturb-seq dataset of Schmidt *et al*. [10]. Primary T cells provide a biologically relevant setting in which genetic perturbations modulate heterogeneous activation states and immune programs [10]. Our benchmark included 57,835 cells spanning 31 annotated cell states (Supplementary Fig. S1), enabling evaluation across a heterogeneous repertoire of immune states. These baselines span the dominant paradigms for predicting post-perturbation single-cell states from control cells and perturbation identity: GEARS [3], a gene-network-guided graph neural network; scGPT [4], a pretrained single-cell foundation model; CPA [12], a compositional perturbation autoencoder; and CellFlow [5], a flow-matching generative model (Supplementary Note 3). We evaluated predictions for held-out perturbations across five cross-validation folds. For each expression-centric baseline, weighted *k*-nearest-neighbor (wKNN) label transfer [21] assigned cell types to predicted expression profiles using a training-derived PCA reference. We aggregated these labels into cell-type proportions for comparison with observed compositions (Methods).

We first compared the observed and predicted cell-state distributions using shared-reference UMAP visualizations. For visualization, each held-out perturbation was evaluated within its fold-matched reference: PCA was fitted on non-held-out training/validation ground-truth cells, UMAP was constructed from those PCA coordinates, and held-out ground truth together with all available model predictions were projected into this shared reference after gene alignment. For the held-out FOXD2 condition, expression-centric models failed to reproduce the observed distribution in the shared UMAP: predicted profiles concentrated in fewer regions than observed profiles (Fig. 1d). The same projection protocol was applied across cross-validation splits for additional held-out perturbations (Supplementary Fig. S2).

**Figure 1.**
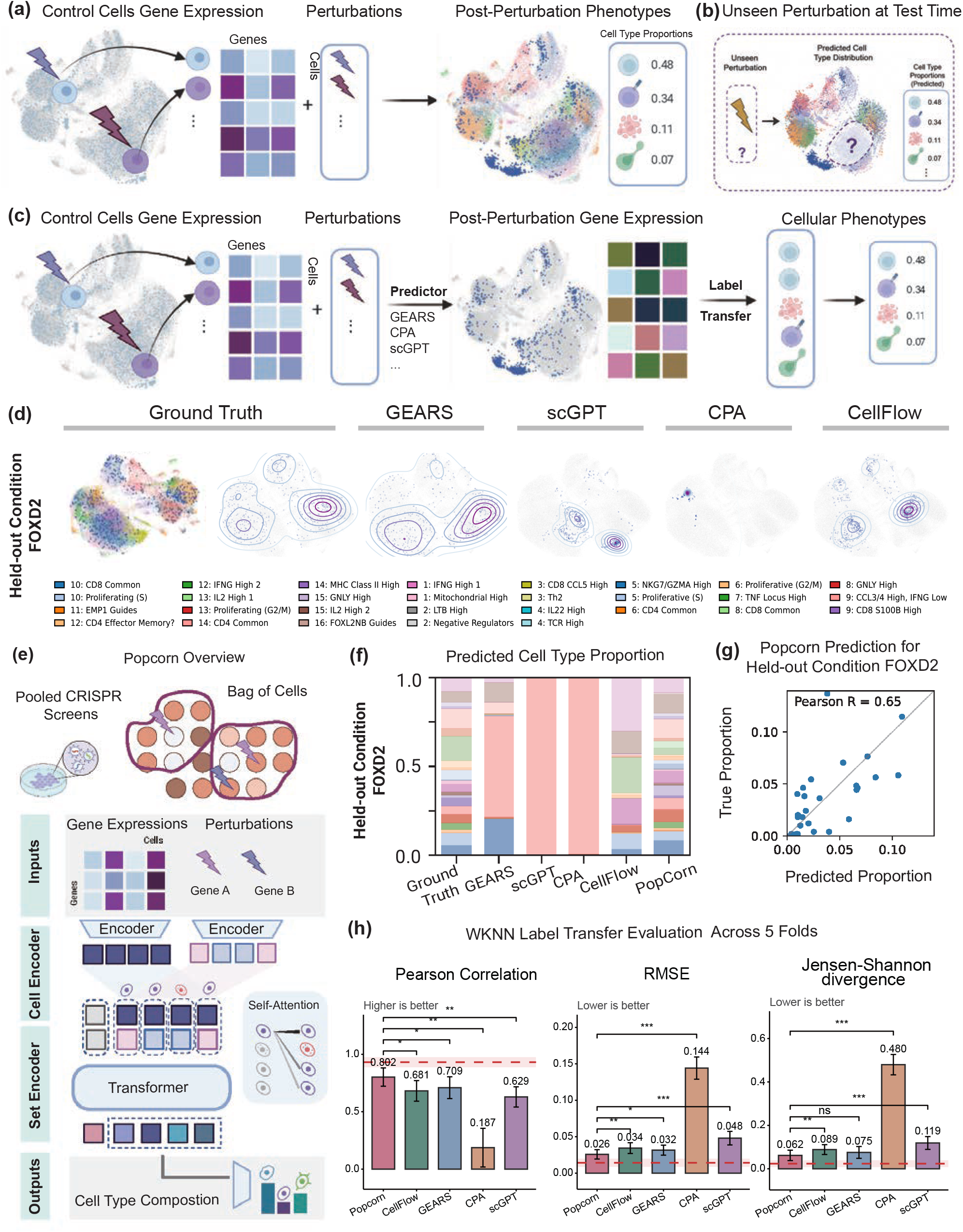
Direct prediction of post-perturbation cell-type composition by Pop-Corn. (a) **Task formulation**. Given baseline (control) single-cell gene expression and a specified genetic perturbation, the objective is to predict the resulting phenotype, defined as the cell-type composition under unseen perturbations. (b) **Generalisation to unseen perturbations**. At test time, Pop-Corn predicts the cell-type composition induced by a perturbation target that was excluded from model training. (c) **Conventional perturbation phenotype prediction pipeline**. Traditional approaches first predict post-perturbation gene expression from control cells and perturbation information, followed by inference of cell-type identities and population proportions through label transfer or downstream annotation. (d) **Ground-truth and predicted post-perturbation expression landscapes**. Single-cell profiles for the held-out FOXD2 perturbation are shown for ground truth and for predictions from GEARS [3], scGPT [4], CPA [12], and CellFlow [5]. Cells are gene-aligned, transformed with a fold-matched PCA model fitted on non-held-out training/validation ground-truth cells, and projected into the corresponding UMAP reference. Colors denote annotated cell types. Expression-centric models may reproduce mean expression but fail to preserve phenotypic distributions. (e) **Pop-Corn framework**. Each perturbation condition is represented as a bag of cells. Perturbations are encoded using ESM protein embeddings of the targeted gene. The model aggregates cell-level representations to learn bag-level phenotypes and is supervised at both cell and bag levels, directly predicting cell-type composition without reconstructing gene expression. (f) **Predicted cell-type compositions**. Stacked bar plots comparing predicted and ground-truth cell-type proportions for the held-out FOXD2 perturbation. Each bar shows the composition across annotated cell types. (g) **Pop-Corn predictions for FOXD2**. Predicted versus observed cell-state proportions for the held-out FOXD2 condition in fold 0. Each point represents one annotated cell state. The displayed Pearson correlation (*r* = 0.65) is calculated across states within this condition. (h) **Benchmark on perturbation phenotype prediction**. Performance on absolute cell-type compositions is summarized across five cross-validation folds using Pearson correlation coefficient (PCC), root mean squared error (RMSE) and Jensen–Shannon divergence (JSD). Pop-Corn achieved a mean PCC of 0.802 across the five folds. The dashed red line is the empirical label-transfer reference obtained from held-out ground-truth expression profiles and is labelled “Test ceiling” in the panel.

We next evaluated predicted cell-type proportions. For the held-out FOXD2 condition, expression-centric pipelines over-assigned cells to dominant states while underrepresenting minority populations (Fig. 1f). We compared predicted and observed compositions using Pearson correlation coefficient (PCC), root mean squared error (RMSE) and Jensen–Shannon divergence (JSD; Fig. 1h). PCC measures linear agreement across cell states, RMSE measures errors in their proportions, and JSD measures distributional dissimilarity. Higher PCC and lower RMSE and JSD indicate better predictions. Together, these results demonstrate that optimizing for expression-level reconstruction does not reliably recover post-perturbation cell-type composition.

### Direct phenotypic mapping improves prediction of post-perturbation population composition

We developed Pop-Corn to predict how a genetic perturbation reshapes a heterogeneous cell population without first reconstructing post-perturbation gene expression (Fig. 1e). Given control single-cell profiles and the identity of a perturbation, Pop-Corn considers the cells jointly and directly predicts the expected cell-type composition under an unseen perturbation. This population-level formulation aligns the model output with the phenotypic outcome of interest, rather than treating composition as a downstream consequence of expression prediction. Details of the model architecture, training objective and control–perturbed cell matching are provided in Methods and Supplementary Note 2 (Supplementary Fig. S5).

Across five cross-validation folds, Pop-Corn achieved the highest mean compositional PCC under wKNN label transfer: 0.802 (GEARS, 0.709; CellFlow, 0.681; scGPT, 0.629; CPA, 0.187). Pop-Corn also had the lowest mean RMSE and JSD (Fig. 1h). These aggregate estimates favored Pop-Corn, although not every pair-wise metric comparison reached statistical significance. Pop-Corn also outperformed the expression-based baselines under cell-type annotation using CellTypist [22] (Supplementary Fig. S4).

For the held-out FOXD2 condition, Pop-Corn more closely recovered the observed cell-type proportions than the expression-centric pipelines (Fig. 1f,g). Additional held-out examples are shown in Supplementary Figs. S2 and S3.

Finally, we evaluated whether Pop-Corn could prioritize perturbations that alter a chosen cell state. We ranked perturbations by their predicted effects on relative cell-state abundance and compared these rankings with those derived from observed compositions. Across screens for increases and decreases, predicted top-five perturbations recovered 70 of 310 observed top-five hits (22.6%; Supplementary Fig. S8). A complementary analysis ranked the cell states most affected by each perturbation (Supplementary Fig. S7).

### Pop-Corn–Spatial predicts niche-level perturbation phenotypes

In spatial perturbation experiments, cells remain embedded within tissue, such that perturbation-associated phenotypes may involve both targeted cells and their local microenvironment. We therefore asked whether Pop-Corn could predict cell-type composition within spatially defined tissue niches.

We evaluated this using the Perturb-Multimodal (Perturb-Multi) dataset of Saunders *et al*. [19], a paired imaging and single-cell sequencing CRISPR screen in mouse liver comprising 5 control and 32 perturbed slices, 202 gene knockouts and 16 annotated cell-state subclusters (Fig. 2a). Because control and perturbed profiles originated from different tissue sections, each perturbed niche lacked a directly observed pre-perturbation counterpart. We therefore used SLAT [23] to establish cross-slice cell correspondences (Fig. 2a). For each perturbation-bearing cell, we defined a 50-*µ*m niche and constructed its pseudo-control from cells within the same radius of the SLAT-matched control anchor (Fig. 2b). Given this aligned reference, its spatial organization and perturbation identity, Pop-Corn–Spatial predicted composition across the 16 annotated subclusters (Fig. 2c). This retrospective evaluation used the observed perturbed tissue to match anchor cells and construct pseudo-control inputs.

**Figure 2.**
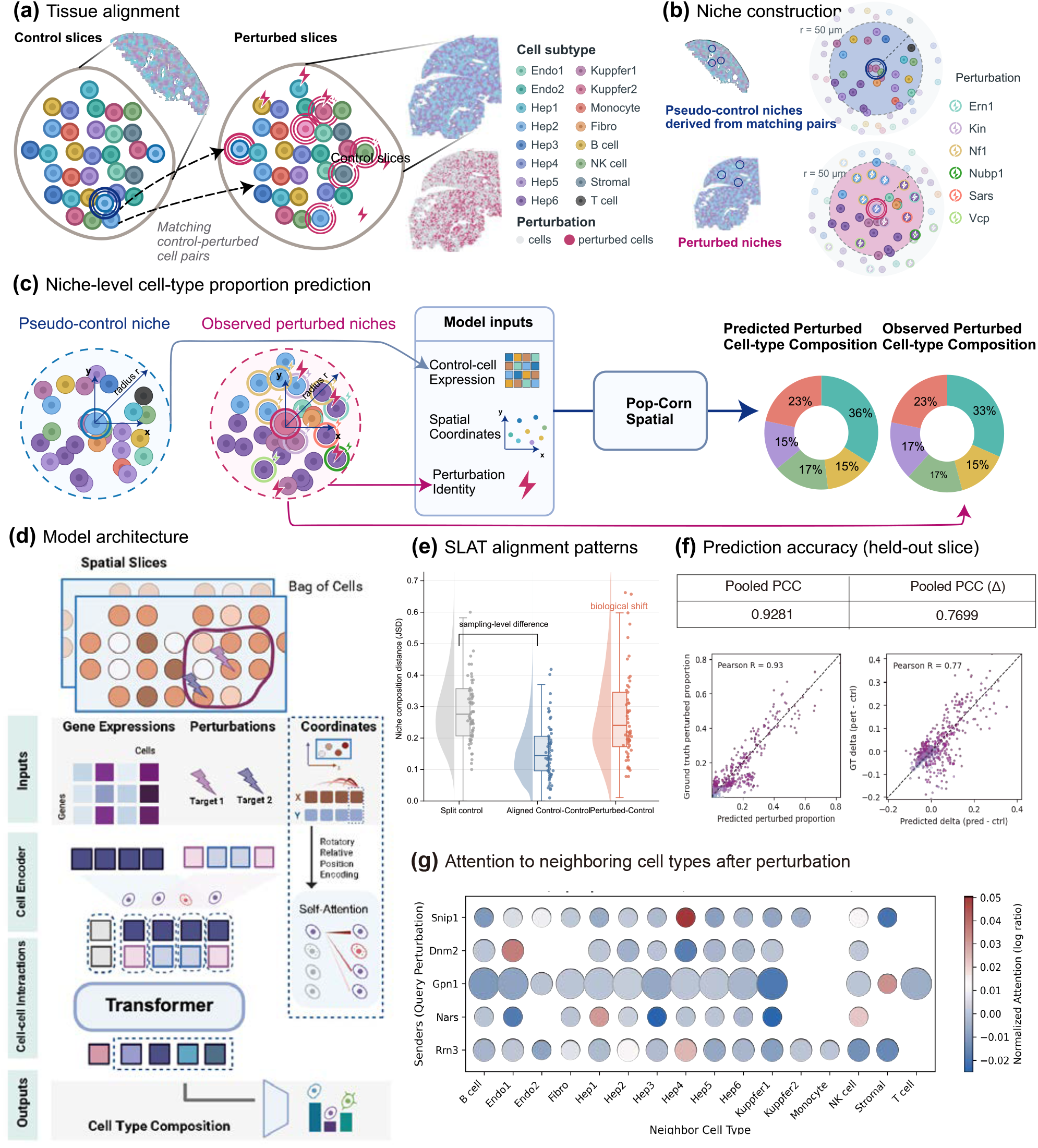
Pop-Corn–Spatial predicts perturbation effects in tissue niches. (a) **Tissue alignment**. SLAT [23] establishes cell correspondences between control and perturbed liver slices from the Perturb-Multimodal dataset [19], retaining cell-subtype and perturbation annotations. (b) **Niche construction**. Cells within 50 *µ*m of a perturbation-bearing anchor cell define a perturbed niche. The matched control counterparts of these cells form its pseudo-control input. (c) **Niche-level cell-type proportion prediction**. Pop-Corn–Spatial receives pseudo-control cell expression, spatial coordinates and perturbation identity, and predicts the corresponding perturbed-niche composition. Observed perturbed-niche composition provides the training target. (d) **Model architecture**. Cell-expression and perturbation embeddings are combined within each spatial bag. Two-dimensional axial rotary positional encoding (2D RoPE) [24] incorporates spatial coordinates into self-attention, and a transformer predicts cell-type composition. (e) **SLAT alignment patterns**. Raincloud plots compare niche-composition Jensen–Shannon divergence (JSD) for split controls, aligned control–control pairs and perturbed–control pairs. Each point represents a niche comparison; lower JSD indicates greater compositional similarity. Comparisons are nested within slice pairs, and differences between perturbed and control slices may also reflect variation between animals or sections. (f) **Prediction accuracy on a held-out slice**. Scatterplots compare predicted and observed proportions for 72 niches and 16 cell states from held-out slice 71. Each point represents a niche–state pair (*n* = 1,152). Pooled Pearson correlation coefficients (PCCs) are 0.9281 for absolute proportions (left) and 0.7699 for changes relative to the aligned pseudo-control (right). Predicted and observed changes subtract the same pseudo-control composition. These correlations summarize this slice only. (g) **Perturbation-conditioned attention**. Rows show five perturbations selected post hoc by their largest positive attention score across cell types; columns show neighbouring cell types. For each query cell, the score is the natural logarithm of the mean attention to a neighbouring cell type divided by the mean attention across all non-self neighbours. Colour shows the median score across contributing query cells: red, above the within-query baseline; blue, below; white, near zero. Point area indicates the number of contributing query-cell records, not neighbour abundance. Blank entries contain no records. These exploratory model readouts do not indicate statistical significance or establish causal cell–cell interactions.

Aligned control–control pairs showed lower niche-composition JSD than split controls and perturbed– control pairs (Fig. 2e). This comparison describes compositional similarity after alignment; it does not establish that a pseudo-control recovers the pre-perturbation state of its matched niche.

Pop-Corn–Spatial incorporates two-dimensional axial rotary positional encoding (2D RoPE [24]) into self-attention (Fig. 2d). RoPE encodes the relative positions of cells along both tissue axes, allowing the model to use their spatial arrangement when predicting niche composition.

We evaluated Pop-Corn–Spatial using leave-one-slice-out cross-validation, excluding all niches from the test slice during training. For the held-out slice shown in Fig. 2f, predicted and observed compositions showed a pooled PCC of 0.9281. Changes relative to the aligned pseudo-control showed a pooled PCC of 0.7699.

Finally, we examined perturbation-conditioned attention for five perturbations selected post hoc by their largest positive attention log ratio across neighbouring cell types (Fig. 2g). Perturbations of *Gpn1, Nars* and *Rrn3*, which affect core transcriptional or translational processes in this screen, showed broad or mixed attention profiles rather than a common cell-type-specific pattern [19]. By contrast, *Dnm2* and *Snip1* contained localized positive deviations for Endo1 and Hep4, respectively. The *Dnm2*–Endo1 pattern was compatible with the established role of DNM2 in endothelial integrin trafficking and morphogenesis [25], whereas the *Snip1*–Hep4 pattern remains an unvalidated candidate. These post-hoc examples nominate gene–cell-type associations for further testing but do not provide independent validation or establish perturbation effects or causal cell–cell interactions.

## Discussion

Pop-Corn shows that perturbation-induced population phenotypes can be predicted directly, without reconstructing the post-perturbation transcriptome. Across held-out perturbations in heterogeneous primary T cells, Pop-Corn recovered cell-type composition more accurately than expression-first pipelines and preserved low-abundance states that those approaches tended to collapse into dominant populations. These results show that accurate transcriptome reconstruction is neither necessary nor sufficient for predicting perturbation-induced changes in cell-type composition. Pop-Corn extends this phenotype-first approach from dissociated cells to intact tissue, where spatial context influences local population structure. Together, these findings establish population composition as a direct, interpretable target for perturbation prediction across both settings.

Pop-Corn fills a gap left open by existing approaches. Neural optimal transport methods such as CellOT model perturbation-induced transitions between cellular states but do not directly optimize annotated cell-state proportions [8]. Distributional generative models such as CellFlow model perturbation-conditioned cell-population distributions, including in developmental and organoid systems [5]. Pop-Corn instead makes compositional change the explicit learning objective. We demonstrate this formulation across held-out perturbations in heterogeneous primary cells and across tissue slices using leave-one-slice-out cross-validation.

Despite the higher predictive performance, a potential limitation of population-level modeling is sampling variability in the observed cell-type proportions. Because only a finite number of cells are measured in each sample or spatial niche, the estimated proportions can vary by chance, particularly for rare cell types and small niches. As a result, some differences between predicted and observed compositions may reflect sampling noise rather than true prediction error, complicating the interpretation of performance metrics.

In the present study, we mitigated sampling variability by aggregating cells at the perturbation or niche level, thereby reducing variance relative to single-cell predictions. Nevertheless, Pop-Corn does not explicitly model sampling uncertainty, and observed proportions are treated as point estimates. Future extensions could incorporate probabilistic models of compositional noise, for example Dirichlet– multinomial or Bayesian hierarchical formulations, to distinguish biological perturbation effects from sampling-induced fluctuations. Uncertainty-aware objectives may further improve robustness, particularly for rare cell populations and sparsely sampled spatial niches.

Finally, Pop-Corn deliberately does not reconstruct gene expression and therefore does not by itself resolve the gene-level mechanisms, including transcription factors and pathways, that drive compositional shifts. This is a design choice rather than a deficiency: because expression accuracy and compositional accuracy are empirically decoupled, reconstructing the full transcriptome is not required for predicting phenotype. Expression-based models remain complementary when gene-level molecular responses are of interest. In this sense, the two approaches address different questions: Pop-Corn predicts what changes in a cell population, whereas expression-centric models can help investigate why. Hybrid architectures that jointly predict composition and expression therefore represent a natural next step, combining phenotype-level prediction with molecular interpretation.

## Methods

### Problem formulation

#### True perturbation-induced phenotypic outcome

Each genetic perturbation *g* shifts the distribution of cellular states in a population. We represent each cell by its log-normalized expression profile **x** ∈ ℝ^*G*^ across *G* genes, and denote the distribution over cell states under perturbation *g* as *P*_*g*_. The phenotypic response of interest is the cell-type composition: the vector of probabilities that a randomly drawn cell belongs to each of *K* annotated types,

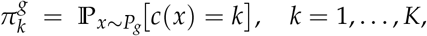

where *c*(*x*) denotes the cell-type label of cell *x*. The composition 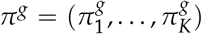 lies on the probability simplex 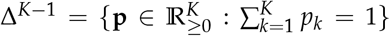, meaning that its entries are non-negative and sum to one. It describes the population as a whole rather than any individual cell.

#### Observed perturbation-induced phenotypic outcome

In practice, perturbation *g* is profiled by sequencing a finite collection of *n*_*g*_ cells, forming an observed bag 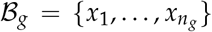 drawn i.i.d. from *P*_*g*_. The directly observable quantity is the empirical composition,

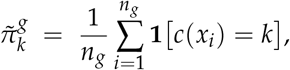

which estimates 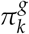with sampling error of order 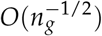. For low-abundance cell types 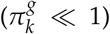 or shallow sequencing (*n*_*g*_ small), this noise is non-negligible and constitutes an irreducible source of label noise in training. Because 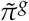 is a property of the bag as a whole rather than of individual cells, predicting it naturally calls for a multiple instance learning (MIL) framework [26].

#### Prediction task

Given a control bag *B*_ctrl_ = {*x*_1_, …, *x*_*n*_} drawn from the unperturbed distribution *P*_0_ and a query perturbation *g*, the goal is to learn a function

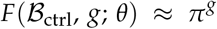

that predicts the true cell-type composition of the perturbed population. Here, *θ* denotes all trainable parameters of Pop-Corn. In training, we supervise against the observable proxy 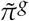; by the law of large numbers, 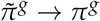 as *n*_*g*_ → ∞, so the two targets coincide in the large-sample regime. In the non-spatial benchmark, perturbations *g* ∉ ℊ _train_ were held out entirely from training. The spatial benchmark used leave-one-slice-out cross-validation, with each test slice and all of its niches excluded from the corresponding training fold.

### Pop-Corn framework

#### Bag construction and sampling strategy

For the Schmidt T cell experiments, each training instance pairs a perturbation identity *g*_*i*_ with control and perturbed bags of equal size *n*. We sampled matched control–perturbed pairs with replacement from the matched pool for *g*_*i*_, retaining both members of each sampled pair:

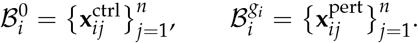

The perturbation identity *g*_*i*_ conditions the control input bag. The composition label is computed from the corresponding sampled perturbed cells:

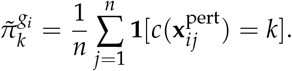

The bag-level composition loss is permutation-invariant and does not itself require cell-to-cell correspon-dence (Supplementary Note 1). We established one-to-one control–perturbed pairings using CINEMA-OT [27] with an exact linear-programming assignment solver to provide cell-level supervision targets (Supplementary Note 2). Conditional on the matched pool, sampling pairs uniformly with replacement yields an unbiased estimate of that pool’s empirical perturbed-cell composition.

#### Bag size

For the non-spatial Schmidt T cell experiments, the effective bag size was determined by bootstrap subsampling rather than by a single fixed integer. For a matched pool containing *N* available control–perturbed cell pairs, each training bag contained *n* = max(1, ⌊0.3*N⌋*) sampled pairs with replace-ment. We generated 50 bootstrap bags during training for each fold. Larger sampling fractions give a better representation of each population but increase computation in the transformer aggregator; the complete run configuration is reported in Supplementary Table S1.

#### Spatial bag construction

Because perturbed and control cells were profiled in separate tissue sections, we used SLAT [23] to establish cell-level correspondences between each perturbed slice and a selected control reference. SLAT combines molecular features with spatial-neighbourhood topology and returns the explicit cross-slice matches required for pseudo-control construction.

Each non-control perturbed cell served as an anchor, and its niche comprised cells within a 50-*µ*m radius. Niches containing fewer than five cells were excluded, and each niche was capped at 1,000 neighbours. The SLAT-matched control counterpart of each perturbed anchor served as the control anchor. Cells within a 50-*µ*m Euclidean radius of this control anchor in the control slice formed the pseudo-control input.The observed composition of the perturbed niche provided the prediction target. In the slice 71 fold shown in Fig. 2f, the matched control reference was Adlib 10. This procedure produced 72 test niches containing 74–229 cells (mean, 145.3). Pseudo-control construction therefore depended on cross-slice anchor matching using the observed perturbed tissue, defining a retrospective prediction task conditional on the aligned reference.After niche construction, coordinates within each slice were min–max scaled to [0, 1]^2^ for input to the 2D RoPE module.

### Multiple instance learning formulation

Because the cell-type composition is a population-level property of the bag rather than of any individual cell, we adopt a *multiple instance learning* (MIL) framework [26, 28, 29], in which the label is attached to the bag as a whole. The model *F* encodes each cell, conditions it on the bag-level perturbation by adding the perturbation representation, aggregates across the bag via a transformer, and decodes a compositional prediction; each component is described below.

### Model architecture

#### Cell encoder

For the Schmidt T cell experiments reported in the main cross-validation benchmark, each cell was represented by a 512-dimensional pretrained scGPT embedding (`X_scgpt_CP_512`), which was projected to the shared hidden dimension *d*_*h*_ by a two-layer feedforward encoder. We compared this representation against alternatives based on the top 100 principal components fitted on the training split and log-normalized counts over the top 5,000 highly variable genes selected on the training split in the cell-encoder ablation (Supplementary Fig. S6). For the spatial Saunders liver experiments, each control cell was represented by its log-normalized expression profile, projected to *d*_*h*_ by the same two-layer feedforward encoder.

#### Perturbation encoder

Each genetic perturbation was represented through the amino acid sequence of its target protein. Gene symbols were mapped to canonical UniProt sequences, which were encoded using the pretrained protein language model ESM-2 (`esm2 t36 3B UR50D`) [30]; residue-level hidden states were mean-pooled across sequence positions and projected to the shared *d*_*h*_-dimensional space via a two-layer feedforward network. Grounding perturbation representations in protein sequence, rather than learning gene embeddings from scratch, confers two practical advantages: it naturally transfers across species (human and mouse paralogs share homologous sequences), and it supports zero-shot prediction for perturbation targets not observed during training.

#### Perturbation conditioning

For each cell *j* in bag ℬ_*i*_, the cell embedding 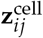 and the perturbation embedding 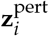 are combined element-wise to form a joint input token,

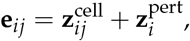

conditioning each cell representation on the shared perturbation identity. Because all cells in the same bag receive the same perturbation embedding, this broadcast also signals to the downstream aggregator which cells belong to the same perturbation context.

#### Transformer aggregator

The set of conditioned tokens (**e**_*i*1_, …, **e**_*in*_) is processed by a bidirectional transformer encoder. In the Schmidt T cell runs, the encoder used one transformer layer with hidden dimension *d*_*h*_ = 64, four attention heads, feedforward dimension 128 and dropout 0.1. This lightweight architecture allows attention across all cells in the sampled bag while keeping the number of trainable parameters small relative to the size of the perturbation screen. For Pop-Corn–Spatial, we adopted a ModernBERT-style encoder with 12 layers, hidden dimension *d*_*h*_ = 768, 12 attention heads, feedforward dimension 3,072 with gated linear units (GLU), pre-norm layer normalization and dropout 0.1.

#### Spatial extension

For spatially-resolved data, each cell additionally carries a 2D tissue coordinate 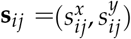. We encode spatial position using 2D axial RoPE [24]: the hidden dimension is split equally between the two spatial axes, each independently rotated, such that the composite rotation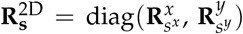 yields attention scores that depend only on the relative displacement **s**^′^ − **s**,

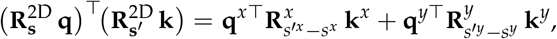

preserving translation equivariance across tissue sections without introducing additional parameters.

#### Compositional decoder

A shared cell-state classification head is applied to each contextualized cell token. The resulting per-cell logits are converted to probabilities by softmax and averaged over valid cells in the bag to obtain the predicted cell-type composition:

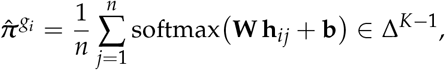

where **h**_*ij*_ is the transformer-contextualized representation of cell *j*, and 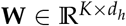 and **b** ∈ ℝ^*K*^ are learned parameters. The softmax normalization ensures that predictions lie on the probability simplex and are directly interpretable as cell-type proportions.

### Learning objective

For the Schmidt T cell experiments, the model is trained with a combination of bag-level and cell-level supervision. In the Schmidt T cell runs, the bag-level term is the mean squared error between the bootstrap composition label 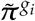 and the predicted composition 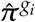. The cell-level term is the cross-entropy between each control cell’s per-cell prediction *ŷ*_*ij*_ and the cell-type label *y*_*ij*_ of its OT-matched perturbed counterpart (Supplementary Note 2), providing a per-cell signal for the transition each control cell would undergo. The two are combined as

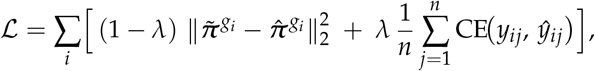

where *λ* ∈ [0, 1] balances the two terms; per-cell logits *ŷ*_*ij*_ are obtained from the shared cell-state head applied to the contextualized cell embeddings. Setting *λ >* 0 activates the cell-level term and substantially improves prediction quality (Supplementary Fig. S5); we use *λ* = 0.5 for the Schmidt T cell experiments. Pop-Corn–Spatial was trained using niche-level composition supervision only. SLAT matching was used to construct pseudo-control inputs, not cell-level supervision targets.

### Inference

At test time, for a held-out perturbation *g* ∉/*G*_*t*rain_, we draw *R* independent control bags of size *n* and average the predictions to reduce sampling variance:

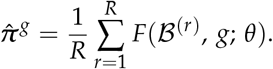

For the Schmidt T cell experiments, we used *R* = 20 bootstrap bags per held-out perturbation. The evaluation target is 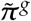 computed from the complete profiled perturbed population *C*_*g*_. To isolate perturbation-induced changes from baseline proportions, we additionally report all metrics on compositional shifts 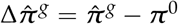, where ***π***^0^ is the control composition.

## Datasets and preprocessing

### T cell CRISPR Perturb-seq datasets

#### Schmidt *et al* [10]

This dataset profiles CRISPR-mediated gene activation (CRISPRa) perturbations in human primary T cells under both stimulated and resting conditions. Our benchmark used 57,835 cells, matching the total across the training/validation and test partitions in each of the five folds (Supplementary Fig. S1c). Raw count matrices were downloaded from the National Institutes of Health Gene Expression Omnibus (GEO; accession GSE190604), and the associated code and data archive was obtained from the versioned Zenodo record [31]. We directly adopted the subcluster annotation provided in the original study, which categorized the T cell population into 31 transcriptionally distinct subclusters. These annotations were derived based on integrated transcriptomic and activation-state signatures curated by the original authors. Cells not annotated in the metadata were excluded in our study. Preprocessing was performed using the `Scanpy` toolkit [32]. Cells with fewer than 500 detected genes and genes expressed in fewer than five cells were filtered out. Expression counts were normalized to a total of 10,000 counts per cell and log_1*p*_-transformed. Highly variable genes (HVGs) were selected using `pp.highly_variable_genes` with the top 5,000 genes retained for downstream analyses.

### Spatial CRISPR Perturb-seq datasets

#### Saunders *et al* [19]

The dataset measures the effects of pooled CRISPR perturbations in situ in mouse liver tissue using the Perturb-Multimodal platform, which integrates in situ imaging and single-cell sequencing readouts. It comprises sgRNAs (identifying the induced genetic perturbation), mRNA gene expression, and the spatial coordination of each cell within tissue slices. The dataset includes both control and CRISPR-perturbed liver tissue slices from mice. The control group comprises five unperturbed (“Adlib”) slices, encompassing 177,392 cells and 13,680 detected genes. The perturbed group consists of 32 slices subjected to pooled CRISPR-Cas9 gene knockout, encompassing 2,203,099 cells and 209 profiled genes corresponding to 202 targeted knockouts. Standard preprocessing procedures were applied, including quality control filtering, normalization, and log_1p_ transformation. Cells were systematically annotated into nine major cell types and further subdivided into sixteen subclusters by domain experts. Spatial coordinates within each slice were subsequently normalized using the max–min method to facilitate downstream analyses and modeling. The processed datasets with fine-grained cell-type and subtype annotations were obtained directly from the authors.

#### Cross-slice matching

Each perturbed slice was compared with all candidate control slices using SLAT. Cells were matched by molecular similarity and spatial proximity after tissue alignment. The control slice with the smallest average spatial distance between matched cells was selected to construct pseudo-controls.

#### SLAT alignment benchmark

To evaluate whether SLAT compared corresponding tissue regions, we analyzed split-control, aligned control–control and perturbed–control comparisons. For each pair, SLAT was applied to align spatial coordinates, and cell-type proportions were compared between aligned regions. The split-control group comprised five within-slice comparisons (10 → 10, 11 → 11, 12 →12, 20→20 and 21→21), yielding 59,381 valid niche-level JSD values. The aligned control–control group comprised ten directed pairs (10 → 11, 11→ 10, 11→ 12, 12 → 11, 10→ 20, 20→ 10, 20→ 21, 21→ 20, 11 → 21 and 21 → 11), yielding 167,559 valid values; 60,000 were retained for plotting. The perturbed–control group yielded 604 valid values from four pairs (80 →12, 90 →12, 91→ 12 and 102→ 12). Candidate pairs 50→ 12 and 52→ 12 yielded no valid records and were not plotted. Each plotted observation represents one valid niche-level JSD rather than a slice-pair mean. These observations are nested within slice pairs and should not be interpreted as independent slice-level replicates. Control–control comparisons measure compositional similarity between aligned regions; they do not independently establish recovery of a perturbed niche’s pre-perturbation state. Residual differences in perturbed–control pairs may include perturbation effects together with animal, section and technical variation. Niche-level composition similarity was quantified by the Jensen–Shannon divergence (JSD) between aligned cell-type proportion vectors; lower values indicate more similar compositions.

### Benchmark on perturbation phenotype prediction tasks

To compare our end-to-end phenotype prediction with baselines, we constructed two-step pipelines in which an expression prediction model generated post-perturbation profiles, followed in the main analysis by wKNN label transfer in a training-derived PCA space. A CellTypist-based annotation pipeline was used as a supplementary sensitivity analysis.

#### Perturbation prediction models

We used four perturbation prediction models to generate post-perturbation expression profiles or cell-state representations from control cells and perturbation targets. Each model was adapted from its official codebase and trained on the same perturbation-level data splits, with target representations and model details provided in Supplementary Note 3.

##### GEARS [3]

GEARS predicts a post-perturbation expression vector for each control-cell input using gene co-expression and Gene Ontology graphs; its original evaluation emphasizes mean differential-expression accuracy. Held-out targets absent from the perturbation graph or excluded during graph preprocessing could not be predicted; these conditions were omitted from GEARS-specific aggregate summaries rather than imputed and were tracked separately in the benchmark outputs.

##### scGPT [4]

scGPT is a transformer foundation model pretrained on single-cell transcriptomes. We fine-tuned it with control-cell expression and perturbation tokens to predict post-perturbation gene expression under a masked mean-squared-error objective, making it an expression-centric comparator.

##### CPA [12]

CPA learns a perturbation-invariant basal cell state and combines it additively with perturbation and covariate representations before decoding post-perturbation expression. In our implementation,perturbation targets were represented by the same pretrained ESM-2 protein embeddings used for Pop-Corn (`esm2_t36_3B_UR50D`), rather than embeddings learned only from training-set perturbation identities. This representation enabled conditioning on targets held out from model training.

##### CellFlow [5]

CellFlow uses conditional flow matching to transport a distribution of control cells to a predicted post-perturbation expression distribution. We conditioned CellFlow on the same pretrained ESM-2 target-protein embeddings used for Pop-Corn (`esm2_t36_3B_UR50D`), permitting predictions for targets held out from model training. CellFlow models population heterogeneity explicitly, but is not trained directly against perturbation-induced cell-type composition.

#### Cell-type label transfer

For the main two-step benchmark, predicted expression profiles from each baseline were projected into a common PCA coordinate system fitted on the labeled non-held-out cells for the corresponding cross-validation fold. Cell-type labels were then transferred from the labeled reference cells to each predicted cell by weighted *k*-nearest-neighbor matching [21] in this PCA space, and predicted cell labels were aggregated within each perturbation condition to estimate cell-type composition. This protocol ensures that all model predictions are evaluated under an identical fold-specific reference projection.

For UMAP visualisation of held-out perturbations, each cross-validation split was processed independently. Within a split, PCA was fit on non-held-out training/validation ground-truth cells using 50 components. A UMAP reference was constructed from these PCA coordinates with 20 nearest neighbours and random seed 42. Held-out ground-truth cells and all available model predictions were aligned to the same gene order, transformed by the same PCA model, and projected into the same UMAP reference using Scanpy ingest.

As a supplementary sensitivity analysis, predicted expression profiles were also passed to CellTypist [22] for cell-type label assignment. CellTypist was trained on the labeled training perturbation conditions (excluding the held-out set) and uses logistic regression classifiers with automated feature selection to assign cell-type labels from normalized gene expression input.

#### Evaluation settings

For all non-spatial perturbation prediction tasks, datasets were split by perturbation condition using a fixed random seed (42). Specifically, 20% of perturbation targets were held out as the test set, while the remaining 80% were used for training and validation. To ensure robustness and reduce variance, we performed 5-fold evaluations, each time rotating the held-out perturbation subset as the test set. For the spatial Perturb-seq evaluation, we used leave-one-slice-out cross-validation. Each evaluated slice was held out in turn, and all niches from that slice were excluded from the corresponding training fold. The slice 71 fold shown in Fig. 2f used batch size 8 and random seed 42.

During the test phase, we used the trained perturbation prediction model to generate perturbed expression profiles or latent embeddings for the held-out perturbations. For the main benchmark, the predicted profiles were projected into the fold-specific reference PCA space and assigned cell-type labels by wKNN transfer from labeled non-held-out cells. For each perturbation condition, we aggregated predicted cell-type assignments to obtain the estimated cell-type composition.

Model performance was evaluated by comparing the predicted and ground-truth cell-type distributions.

#### Assessment metrics

For each held-out perturbation, we compared the predicted composition with the empirical composition of the profiled perturbed cells. Pearson correlation coefficient (PCC) measured linear agreement across cell types, root mean squared error (RMSE) measured coordinate-wise error and Jensen–Shannon divergence (JSD) measured distributional dissimilarity. Higher PCC and lower RMSE or JSD indicate better performance. The aggregate benchmark and ablation figures additionally report mean squared error (MSE), Spearman correlation coefficient (SCC) or the coefficient of determination (*R*^2^), as indicated. PCC, MSE, RMSE, SCC and *R*^2^ were evaluated on either absolute compositions or perturbation-induced shifts relative to the empirical composition of the complete control population; JSD was evaluated only on absolute compositions. For the non-spatial benchmark, metrics were computed for each held-out condition and summarised across the five cross-validation folds. As a label-transfer reference, we applied the same annotation pipeline to held-out ground-truth expression profiles; this reference is shown as a dashed line in the benchmark figures. Formal definitions are provided in Supplementary Note 4. For Fig. 2f, predicted and observed composition matrices for 72 niches from held-out slice 71 were each flattened across 16 cell states. Pearson correlation was calculated between the resulting vectors of 1,152 niche–state entries. For each niche, predicted and observed changes were defined as 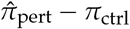 and 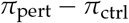, respectively. Both used the same matched pseudo-control composition, and their correlation was calculated using the same flattening procedure. These pooled correlations were descriptive summaries of the displayed slice; niche–state entries were not treated as independent biological replicates.

#### Spatial attention analysis

For the attention analysis shown in Fig. 2g, we extracted self-attention from the final transformer layer (layer 11 of 12) and averaged the weights across all 12 heads. Each query was a perturbed cell, and its keys were all other cell tokens in the same niche; self-attention entries were excluded. We used all non-self neighbours without distance stratification. For query cell *q* and neighbouring cell type *c*, the normalized attention score was

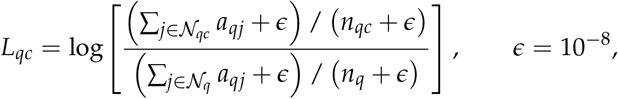

where *N*_*qc*_ contains the non-self neighbours of type *c, N*_*q*_ contains all non-self neighbours, and *a*_*qj*_ is the head-averaged attention from query *q* to key *j*. Thus, the denominator is the mean attention over the same query’s neighbourhood, rather than a control-condition baseline. Scores were not further standardized. Within each perturbation–neighbour-type combination, we summarized *L*_*qc*_ across query cells by the median. Dot colour encodes this median. Dot area was set to 120*m* + 20, where *m* is the number of contributing query-cell records; it does not represent neighbour abundance or attention mass. Combinations without records were omitted. The five displayed perturbations were selected post hoc by ranking each perturbation according to its largest positive median score across neighbouring cell types and retaining the top five.

#### Retrospective virtual screening

We evaluated two ranking tasks within the held-out perturbation set of each cross-validation fold. For perturbation-to-state screening, cell states were ranked by the absolute predicted or observed log_2_ fold-change in relative abundance from control. For state-to-perturbation screening, held-out perturbations were ranked separately for increases and decreases using signed log_2_ fold-changes for each cell state. Within each fold, we counted the overlap between predicted and observed top-*k* sets, using *k* = 15 for perturbation-to-state screening and *k* = 5 for state-to-perturbation screening. We then aggregated results across all folds, reporting recovery as the total overlap divided by the total number of observed top-ranked entries.

## Acknowledgements and funding

This work was supported by the Swiss National Science Foundation Grant No 192128 to M.R.M.. This work was also supported by the Whitehead Institute and the Whitehead Innovation Initiative. N.S. is supported by the Whitehead Institute Fellows Program, the Whitehead Innovation Initiative, and the NIH Director’s Early Independence Award (DP5OD042638).

## Author Contributions

J.C., Y.C., Y.S., N.S., and M.R.M. conceptualized the project and formulated the population-level perturbation phenotype prediction problem. J.C., Y.C., and Y.S. designed the Pop-Corn framework, including the multiple-instance learning formulation, the transformer-based aggregator, the perturbation-conditioning scheme, and the combined bag- and cell-level supervision objective. J.C. extended the framework to spatial Perturb-seq data, implemented all models, curated and preprocessed the Perturb-seq datasets, designed and conducted all benchmarking and ablation experiments, performed the analyses, generated all figures, and drafted the manuscript. M.R.M. and N.S. supervised the project. M.R.M. advised on methodology and presentation. N.S. supervised the biological applications on Perturb-seq data and spatial perturbation data. All authors discussed the results, contributed to manuscript revision, and approved the final version.

## Data Availability

The primary human T cell CRISPRa Perturb-seq data analysed in this study are publicly available from the Gene Expression Omnibus under accession GSE190604, with the associated code and data archive available from the versioned Zenodo record [31]. The Perturb-Multimodal spatial CRISPR dataset is available from the Gene Expression Omnibus under accession GSE275483 and from Hugging Face at https://huggingface.co/datasets/xingjiepan/PerturbMulti. Processed data supporting the main and supplementary analyses,including processed AnnData objects, cell and perturbation metadata, cross-validation splits, figure source-data tables and derived prediction outputs, are available in the Pop-Corn processed-data repository on Hugging Face at https://huggingface.co/datasets/GabbyKoki/Pop-Corn-processed-data.

## Code Availability

The source code implementing Pop-Corn is available at https://github.com/GabbyKoki/pop.

