## Supplementary Information for "Pop-Corn: Predicting Perturbation Phenotype Effects Across Single-Cell and Spatial Contexts"

### Pop-Corn: Supplementary Information

### Supplementary Notes

#### Supplementary Note 1: Statistical Analysis of Phenotypic Composition Estimation

##### Setup and notation

Each perturbation $g$ induces a probability distribution $P_g$ over cell states $\mathcal{X} \subseteq \mathbb{R}^G$. Let $c : \mathcal{X} \to [K]$ be a cell-type annotation function. The *true phenotype* of perturbation $g$ is the vector of cell-type probabilities under $P_g$:

$$\pi_k^g = \mathbb{P}_{x \sim P_g}[c(x) = k], \quad k = 1, \ldots, K, \tag{S1}$$

where $\pi^g = (\pi_1^g, \ldots, \pi_K^g) \in \Delta^{K-1}$. This is a *population-level* quantity: it cannot be measured directly, only estimated from finite samples.

##### Finite-sample estimation of composition

Suppose we observe $n_g$ cells i.i.d. from $P_g$, each independently labeled as one of $K$ types. The empirical composition is

$$\tilde{\pi}_k^g = \frac{1}{n_g} \sum_{i=1}^{n_g} \mathbf{1}[c(x_i) = k], \quad k = 1, \ldots, K, \tag{S2}$$

and the count vector $n_g \tilde{\pi}^g \sim \mathrm{Mult}(n_g, \pi^g)$.

**Proposition 1** (Concentration). *For any fixed cell type $k$ and $\delta \in (0, 1)$, with probability at least $1 - \delta$,*

$$|\tilde{\pi}_k^g - \pi_k^g| \leq \sqrt{\frac{\log(2/\delta)}{2n_g}}, \tag{S3}$$

*by Hoeffding's inequality applied to i.i.d. Bernoulli($\pi_k^g$) variables. As a concrete example: with $n_g = 1000$ cells and $\delta = 0.05$, the per-type absolute error is at most $\approx 4.3\%$ with 95% probability.*

#### Consistency of training on empirical labels

Pop-Corn is trained against empirical cell-type compositions rather than unobserved population compositions. In the Schmidt T cell runs reported here, the bag-level term is mean-squared error between the model prediction $\hat{\pi}^g = F(\mathcal{B}_{\text{ctrl}}, g; \theta)$ and the observed label $\tilde{\pi}^g$:

$$\mathcal{L}_{\text{bag}}(\theta) \;=\; \mathbb{E}_g\Big[\|\tilde{\pi}^g - \hat{\pi}^g\|_2^2\Big]. \tag{S4}$$

Since $\mathbb{E}[\tilde{\pi}^g] = \pi^g$, the true target and the observable label agree in expectation. A standard bias-variance decomposition gives:

$$\mathbb{E}_g\Big[\|\tilde{\pi}^g - \hat{\pi}^g\|_2^2\Big] \;=\; \mathbb{E}_g\Big[\|\pi^g - \hat{\pi}^g\|_2^2\Big] \;+\; \underbrace{\mathbb{E}_g\Big[\|\tilde{\pi}^g - \pi^g\|_2^2\Big]}_{\text{irreducible sampling noise}}\;, \tag{S5}$$

where the second term does not depend on $\theta$ and vanishes as $n_g \to \infty$. Minimizing $\mathcal{L}_{\text{bag}}(\theta)$ over $\theta$ is therefore equivalent to minimizing the population-level squared-error objective up to this irreducible constant. In finite samples, the label noise introduces variance whose magnitude is bounded by the concentration result above.

#### Alternative geometries for compositional prediction

Although the reported Schmidt T cell runs used MSE for the bag-level composition term, KL divergence provides a useful alternative geometry when rare cell types should be weighted more strongly. KL penalizes prediction errors on rare cell types proportionally to $1/\hat{\pi}^g_k$: a 1% absolute error on a 2% cell type is penalized 25× more than the same error on a 50% cell type. Concretely,

$$D_{\text{KL}}(\pi^g \| \hat{\pi}^g) = \sum_{k=1}^{K} \pi^g_k \log \frac{\pi^g_k}{\hat{\pi}^g_k} \;\approx\; \frac{1}{2}\sum_k \frac{(\pi^g_k - \hat{\pi}^g_k)^2}{\hat{\pi}^g_k} \quad (p \approx q), \tag{S6}$$

showing the effective weight $1/\hat{\pi}^g_k$ per cell type. Beyond this geometric property, $-\log \hat{\pi}^g_k$ is a *proper scoring rule*: the unique minimizer of $\mathbb{E}_{x\sim P_g}[-\log \hat{\pi}^g_{c(x)}]$ over all $\hat{\pi}^g \in \Delta^{K-1}$ is $\hat{\pi}^g = \pi^g$. We therefore treat KL-style objectives as a principled extension for settings where rare-state sensitivity is prioritized, while reporting the actual loss used for the Schmidt runs in Supplementary Table S1.

### Supplementary Note 2: Control–Perturbed Cell Matching via Optimal Transport

Pop-Corn's cell-level supervision term assigns a cross-entropy target to each individual control cell in the input bag, requiring a correspondence between control cells and the perturbed cell types they would transition to. In standard Perturb-seq experiments, control and perturbed cells are profiled separately with no lineage tracing, so this correspondence must be inferred. We use CINEMA-OT [1] to establish it: OT finds a coupling that minimizes total transport cost between the control and perturbed populations in a shared PCA embedding space (top 30 principal components), where distances better reflect transcriptomic similarity than raw expression.

**Solver choice: exact assignment over Sinkhorn.** Let $\{x_i^{\text{ctrl}}\}_{i=1}^n$ and $\{x_j^{\text{pert}}\}_{j=1}^n$ denote the control and perturbed cells embedded in the shared PCA space, both equipped with uniform marginals $a = b = \frac{1}{n}\mathbf{1}_n$, and let $C \in \mathbb{R}^{n\times n}$ with $C_{ij} = \|x_i^{\text{ctrl}} - x_j^{\text{pert}}\|_2^2$ be the squared-Euclidean cost matrix. Writing $\Pi(a,b) = \{T \in \mathbb{R}^{n\times n}_{\geq 0} : T\mathbf{1}_n = a,\ T^\top \mathbf{1}_n = b\}$ for the transport polytope, the Kantorovich problem is

$$T^\star \;=\; \arg\min_{T\in\Pi(a,b)} \langle T, C\rangle. \tag{S7}$$

The Sinkhorn algorithm [2] replaces this with an entropy-regularized objective of strength $\varepsilon > 0$,

$$T^\varepsilon \;=\; \arg\min_{T\in\Pi(a,b)} \langle T, C\rangle \;-\; \varepsilon\, H(T), \qquad H(T) \;=\; -\sum_{i,j} T_{ij}\big(\log T_{ij} - 1\big), \tag{S8}$$

whose optimum admits the closed form $T^\varepsilon_{ij} = u_i \exp(-C_{ij}/\varepsilon)\, v_j$ for positive scaling vectors $u, v \in \mathbb{R}^n_{>0}$ recovered by iterated row/column normalization. Since $\exp(-C_{ij}/\varepsilon) > 0$ for every $(i,j)$, the optimal $T^\varepsilon$ has *strictly positive* entries everywhere: each control cell is fractionally matched to all $n$ perturbed cells.

We instead solve the unregularized LP ($\varepsilon = 0$) directly. By the Birkhoff–von Neumann theorem, the vertices of $\Pi(\frac{1}{n}\mathbf{1}_n, \frac{1}{n}\mathbf{1}_n)$ are exactly the scaled permutation matrices $\{\frac{1}{n}P_\sigma : \sigma \in S_n\}$, where $S_n$ is the symmetric group and $(P_\sigma)_{ij} = \mathbf{1}[\sigma(i) = j]$. Because the LP attains its optimum at a vertex, a vertex-returning solver (network simplex / Hungarian algorithm) returns

$$T^\star = \frac{1}{n} P_{\sigma^\star}, \qquad \sigma^\star = \arg\min_{\sigma \in S_n} \sum_{i=1}^{n} C_{i,\sigma(i)}, \tag{S9}$$

which is the classical assignment problem. Each row of $T^\star$ has exactly one nonzero entry: control cell $i$ is matched to the unique perturbed cell $\sigma^\star(i)$. Because cell-level supervision requires a single discrete target per cell, this exact one-to-one matching provides unambiguous per-cell training targets, whereas the dense Sinkhorn coupling would leave the target a convex combination over many candidate perturbed cells.

**OT is only needed under cell-level supervision.** With bag-only supervision ($\lambda = 0$), OT and random bag construction are empirically indistinguishable (Supplementary Fig. S5), because the bag-level composition loss depends only on aggregate cell-type proportions, which are permutation-invariant. OT matching becomes beneficial once cell-level supervision is added ($\lambda > 0$), confirming that its role is to supply meaningful per-cell targets rather than to improve the population-level representation.

#### Supplementary Note 3: Perturbation prediction baseline models

The two-step benchmark compared Pop-Corn with four models that first predict post-perturbation expression profiles or cell-state representations. All models were adapted from their official codebases, trained on the same perturbation-level splits and followed by the common cell-type label-transfer procedure described in the Methods.

**GEARS [3].** GEARS (graph-enhanced gene activation and repression simulator) predicts a post-perturbation expression vector for each control-cell input. Each gene has two learnable embeddings: one refined by a graph neural network over a gene co-expression graph and another by a graph neural network over a Gene Ontology–derived perturbation-similarity graph. Multi-gene perturbations are represented by summing their perturbation embeddings; the combined gene and perturbation representations pass through a cross-gene layer and gene-specific decoders. Training uses an autofocus loss that up-weights large errors together with a direction-aware term penalizing sign mismatches relative to control. This graph structure supports extrapolation to unseen single or combinatorial perturbations when their targets are represented in the graph, and the original evaluation emphasizes mean differential-expression accuracy. Held-out targets absent from the perturbation graph or excluded during graph preprocessing could not be predicted and were omitted from GEARS-specific aggregate summaries rather than imputed.

**scGPT [4].** scGPT is a transformer-based generative foundation model pretrained on more than 33 million human single-cell transcriptomes from the CELLxGENE corpus. Each cell is encoded using gene-identity, binned expression-value and condition tokens, and pretraining uses a masked generative objective to predict held-out gene expression. For perturbation prediction, we fine-tuned scGPT on control and post-perturbation data: the model receives a control-cell expression profile together with tokens identifying the perturbed gene or genes and regresses post-perturbation expression under a masked mean-squared-error loss. Its objective is therefore expression-centric and oriented towards per-gene transcriptional reconstruction.

**CPA [5].** CPA (Compositional Perturbation Autoencoder) learns a disentangled latent representation of perturbation responses. Its encoder maps each cell to a basal state from which perturbation and covariate information is removed adversarially. A perturbed state is then formed by adding perturbation and covariate representations to the basal state, and a decoder maps this state back to gene expression. CPA is trained using an expression-reconstruction objective together with adversarial classification losses and thus serves as an expression-centric comparator. For this benchmark, each target protein was represented by the same 2,560-dimensional, mean-pooled ESM-2 embedding used for Pop-Corn (`esm2_t36_3B_UR50D`). Supplying this sequence-derived representation instead of relying only on a categorical embedding learned for training-set perturbations allowed CPA to condition on held-out targets.

**CellFlow [6].** CellFlow is a generative framework based on conditional flow matching that transports a distribution of control cells to a post-perturbation expression distribution. It fits a conditional velocity field along a probability path between the two distributions and can use optimal-transport couplings to pair sampled source and target cells, producing straighter paths for flow matching. Training regresses the predicted velocity field against the prescribed probability path. For this benchmark, CellFlow was conditioned on the same 2,560-dimensional, mean-pooled ESM-2 target-protein embeddings used for Pop-Corn (`esm2_t36_3B_UR50D`). At inference, the embedding of a held-out target was supplied with the control-cell population to generate post-perturbation expression profiles. CellFlow therefore models population-level heterogeneity explicitly, but it is not optimized directly for perturbation-induced cell-type composition.

#### Supplementary Note 4: Assessment metric definitions

For a held-out perturbation, let $\hat{\mathbf{p}}, \mathbf{p}^* \in \Delta^{K-1}$ denote the predicted and empirical cell-type compositions. For metrics reported on perturbation-induced changes, we instead use $\Delta\hat{\mathbf{p}} = \hat{\mathbf{p}} - \mathbf{p}^0$ and $\Delta\mathbf{p}^* = \mathbf{p}^* - \mathbf{p}^0$, where $\mathbf{p}^0$ is the empirical composition of the complete control population. In the definitions below, $(\mathbf{u}, \mathbf{v})$ denotes either the absolute or delta pair, as specified for each analysis.

**Pearson and Spearman correlation coefficients.** Pearson correlation coefficient (PCC) measures linear agreement across the $K$ cell-type entries:

$$\mathrm{PCC}(\mathbf{u}, \mathbf{v}) = \frac{\sum_{k=1}^{K}(u_k - \bar{u})(v_k - \bar{v})}{\sqrt{\sum_{k=1}^{K}(u_k - \bar{u})^2 \sum_{k=1}^{K}(v_k - \bar{v})^2}}, \qquad \bar{u} = K^{-1}\sum_k u_k, \quad \bar{v} = K^{-1}\sum_k v_k. \tag{S10}$$

Spearman correlation coefficient (SCC) is the PCC between the within-vector ranks of $\mathbf{u}$ and $\mathbf{v}$ and measures monotone agreement. Higher PCC or SCC indicates better agreement.

**Squared-error metrics and coefficient of determination.**

$$\mathrm{MSE}(\mathbf{u}, \mathbf{v}) = \frac{1}{K}\sum_{k=1}^{K}(u_k - v_k)^2, \qquad \mathrm{RMSE}(\mathbf{u}, \mathbf{v}) = \sqrt{\mathrm{MSE}(\mathbf{u}, \mathbf{v})}. \tag{S11}$$

Lower MSE or RMSE indicates smaller coordinate-wise error. The coefficient of determination is

$$R^2(\mathbf{u}, \mathbf{v}) = 1 - \frac{\sum_{k=1}^{K}(u_k - v_k)^2}{\sum_{k=1}^{K}(v_k - \bar{v})^2}. \tag{S12}$$

An $R^2$ of 1 indicates an exact match, whereas values below zero indicate performance worse than predicting the mean of $\mathbf{v}$.

**Jensen–Shannon divergence.** JSD was evaluated only for absolute compositions. With $\mathbf{m} = (\hat{\mathbf{p}} + \mathbf{p}^*)/2$,

$$\mathrm{JSD}(\hat{\mathbf{p}}, \mathbf{p}^*) = \frac{1}{2}D_{\mathrm{KL}}(\hat{\mathbf{p}} \| \mathbf{m}) + \frac{1}{2}D_{\mathrm{KL}}(\mathbf{p}^* \| \mathbf{m}), \tag{S13}$$

where $D_{\mathrm{KL}}(\mathbf{a} \| \mathbf{b}) = \sum_k a_k \log(a_k/b_k)$ with $0 \log 0 = 0$. In numerical evaluation, $10^{-8}$ was added to each entry before the composition vectors were renormalized. Using natural logarithms, JSD lies between 0 and $\log 2$; lower values indicate more similar compositions.

### Supplementary Figures

**(a)**

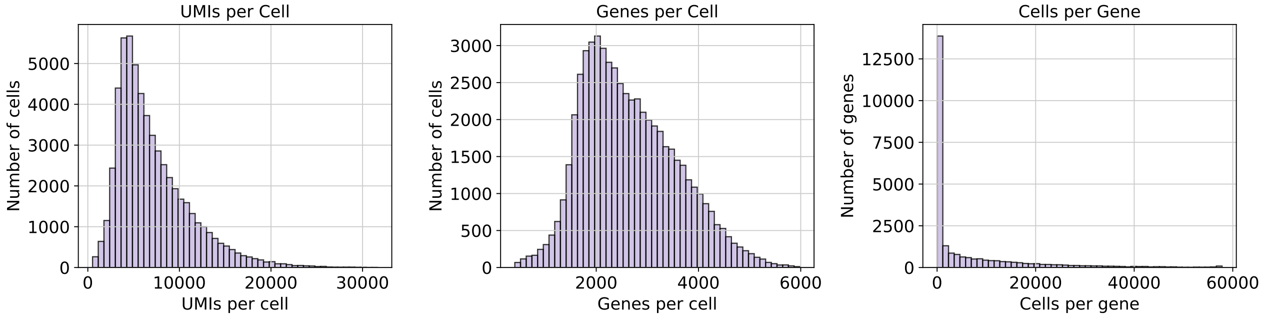


**(b)**

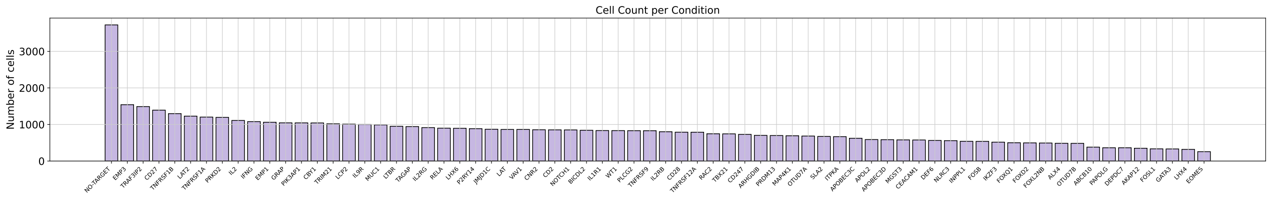


**Fig. S1**: **T cell Perturb-seq dataset overview.** (a) Quality control metric distributions across all profiled cells, including number of detected genes and total UMI counts per cell, after filtering. (b) Number of cells profiled per perturbation condition across all 68 CRISPRa targets. (c) Train/validation/test split by perturbation condition (left) and by cell count (right). (d) Heatmap of cell-type cluster composition per perturbation condition, showing the fraction of cells assigned to each of the 31 transcriptionally distinct subclusters.

**(c)**

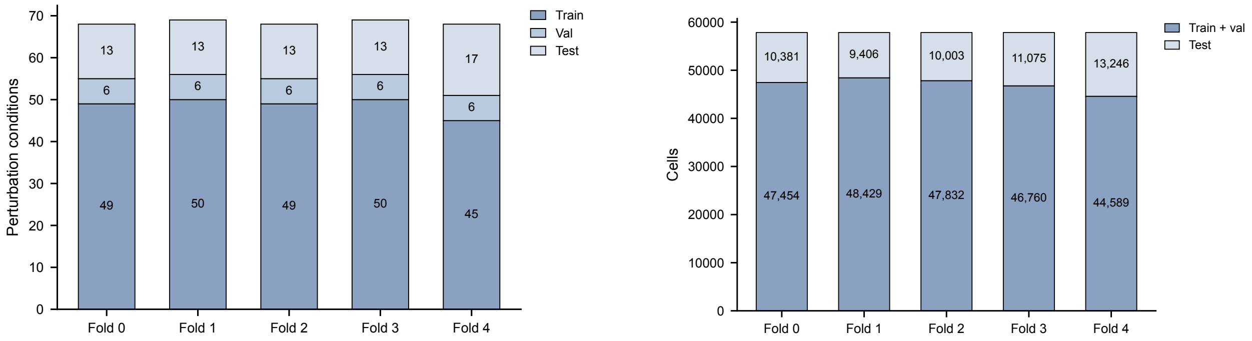


**Fig. S1**: **T cell Perturb-seq dataset overview** (continued). (c) Train/validation/test split by perturbation condition (left) and by cell count (right).

**(d)**

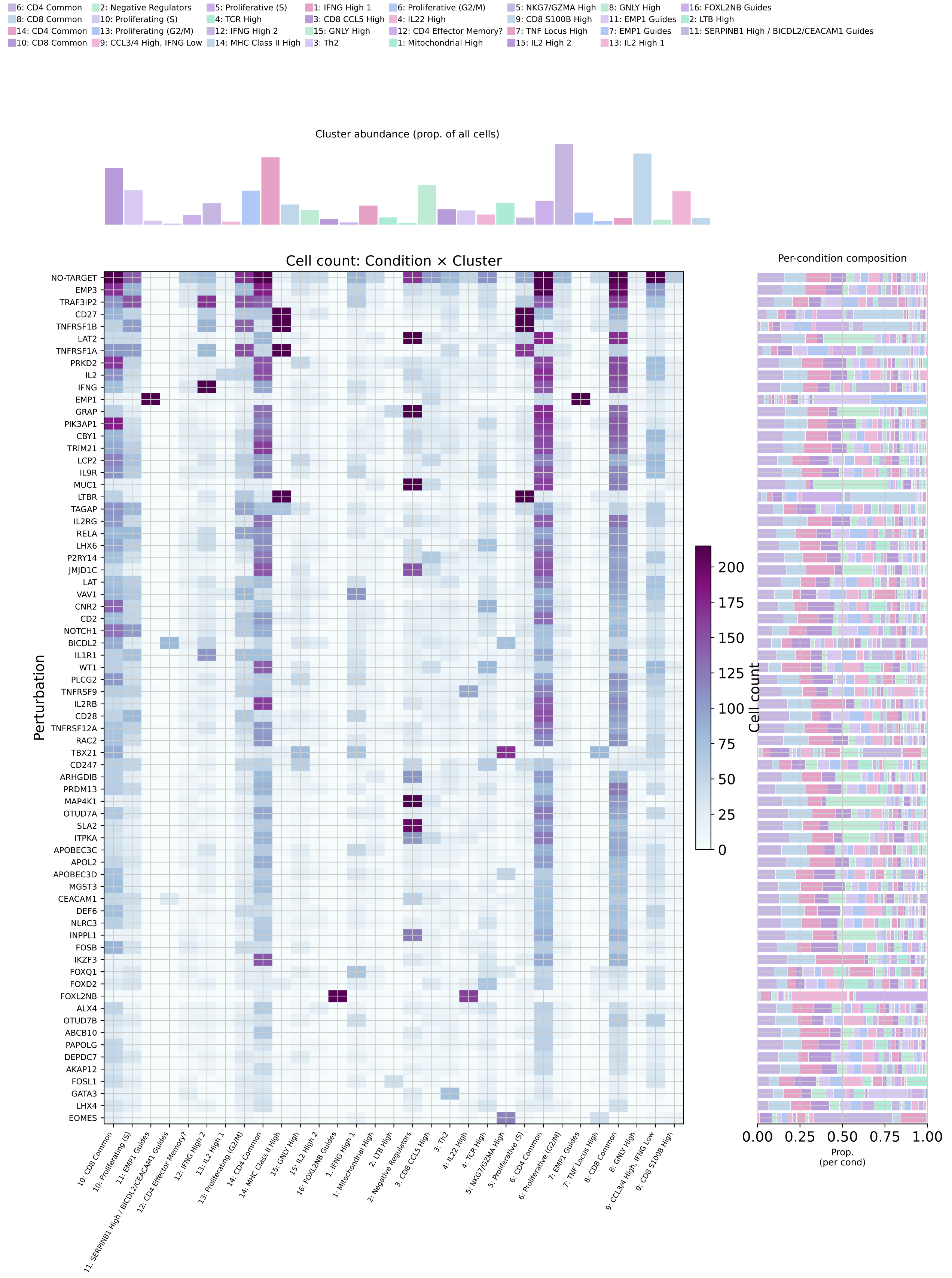


**Fig. S1**: **T cell Perturb-seq dataset overview** (continued). (d) Heatmap of cell-type cluster composition per perturbation condition, showing the fraction of cells assigned to each of the 31 transcriptionally distinct subclusters.

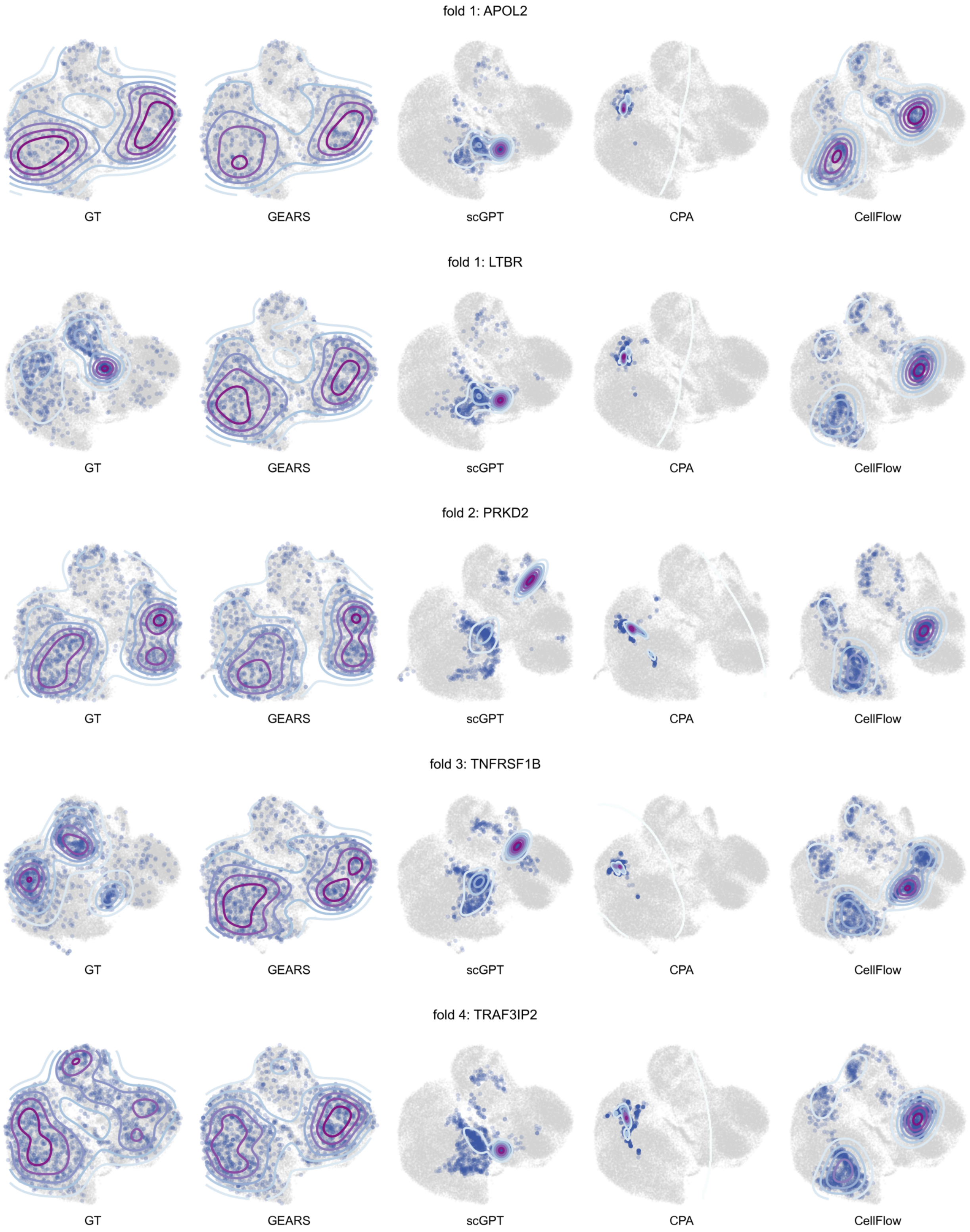


**Fig. S2**: **Additional held-out perturbation landscapes under a shared reference projection.** Five held-out perturbations selected across cross-validation folds (APOL2, LTBR, PRKD2, TNFRSF1B and TRAF3IP2) are shown for ground truth and predictions from GEARS, scGPT, CPA and CellFlow. For each row, PCA and UMAP were fitted only on the corresponding fold's non-held-out training/validation ground-truth cells; held-out ground-truth cells and all available model predictions were gene-aligned, transformed by the same PCA model and projected into the same UMAP reference using Scanpy ingest. These additional examples support the representative visualization in Fig. 1d and show that the observed compression of predicted cellular landscapes is not specific to a single perturbation.

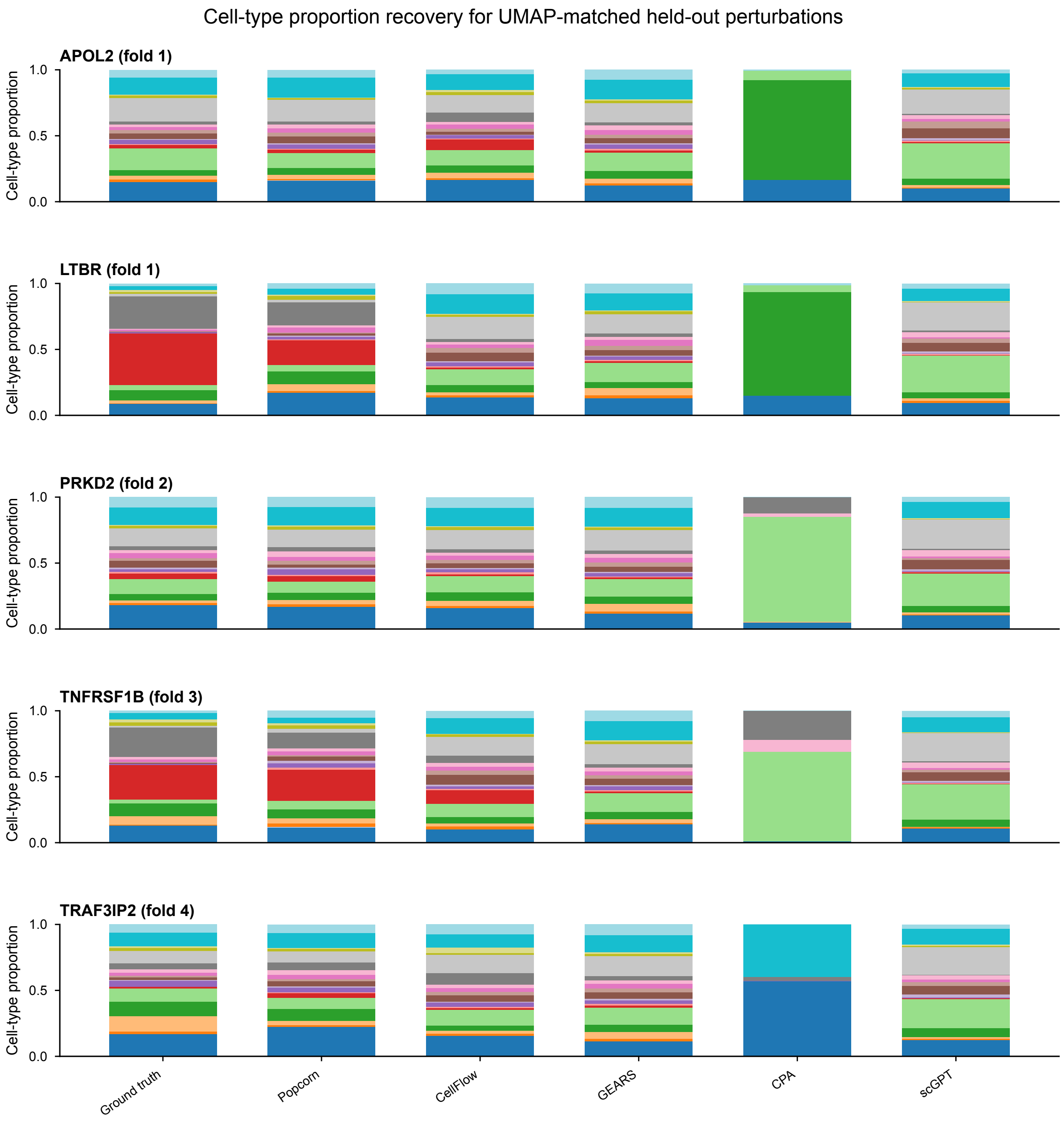


**Fig. S3**: **Cell-type proportion recovery for UMAP-matched held-out perturbations.** Stacked bars show ground-truth cell-type proportions and the corresponding predictions from Pop-Corn and expression-centric baselines for the same five held-out perturbations shown in Supplementary Fig. S2 (APOL2, LTBR, PRKD2, TNFRSF1B and TRAF3IP2). Baseline compositions were inferred from predicted expression profiles using the main weighted *k*-nearest-neighbor label-transfer benchmark; Pop-Corn directly predicts the cell-type proportion vector.

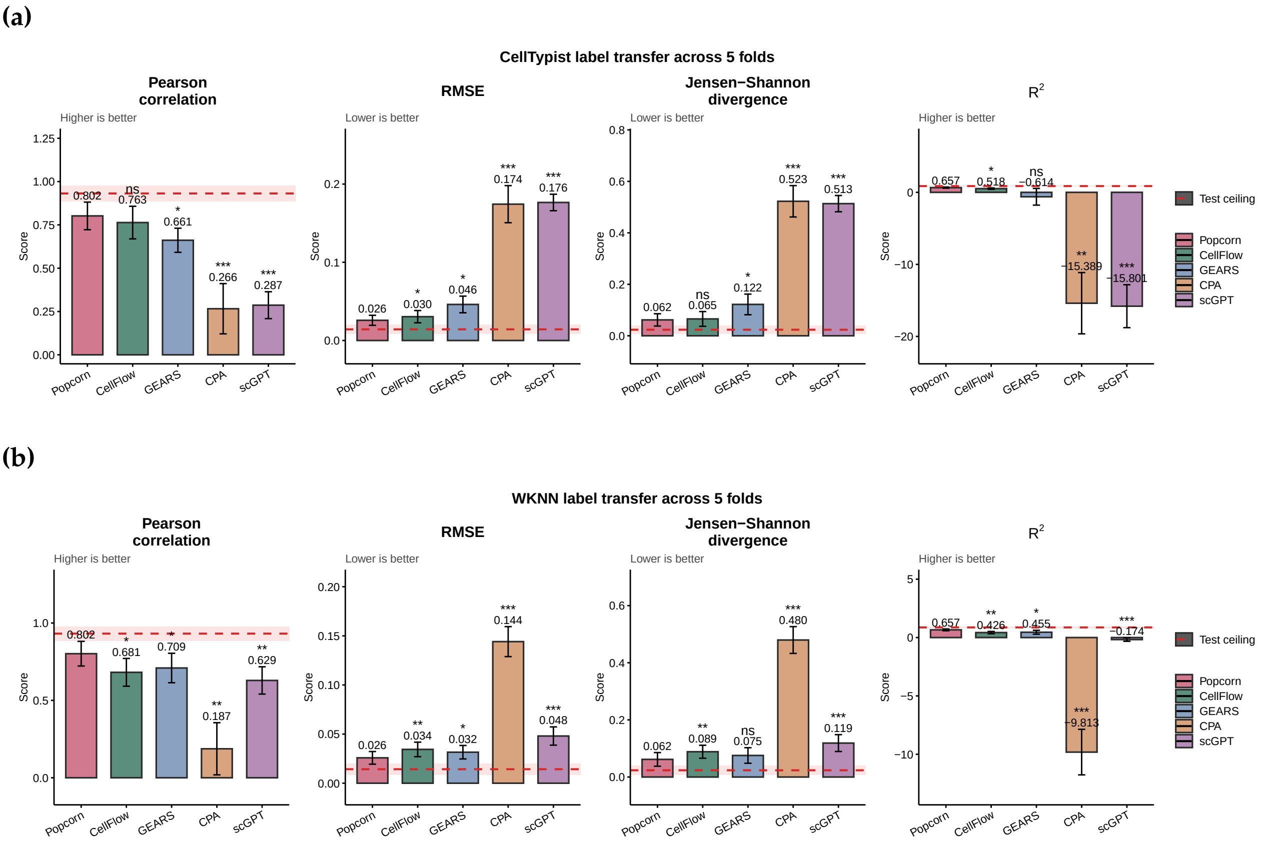


**Fig. S4**: **Aggregate phenotype prediction performance across five-fold cross-validation.** Mean performance on absolute cell-type compositions over five cross-validation folds comparing Pop-Corn against expression-centric two-step pipelines (GEARS, scGPT, CPA, CellFlow). The panels report PCC, RMSE, JSD and $R^2$, with significance markers indicating paired comparisons against Pop-Corn. The dashed red line, labelled "Test ceiling", is the empirical reference obtained by applying the corresponding label-transfer procedure to held-out ground-truth expression profiles. (a) Supplementary sensitivity analysis using Cell-Typist for label transfer. (b) Main benchmark using weighted $k$-nearest-neighbor (wKNN) label transfer in a training-derived PCA space. Pop-Corn outperforms all expression-centric baselines under both label-transfer methods, demonstrating that the advantage is not specific to a particular label-transfer choice.

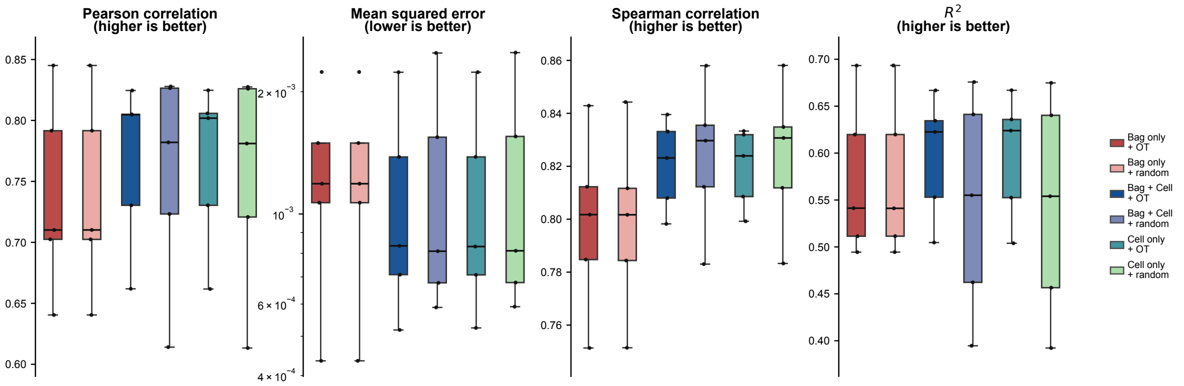


**Fig. S5**: **Cell-level supervision boosts performance, and OT matching further improves it by providing meaningful per-cell training targets.** Five-fold ablation crossing bag-level and cell-level supervision with OT-matched versus randomly sampled control bags. For pure bag-level supervision ($\lambda = 0$), OT has negligible effect, indicating that aggregate phenotype prediction does not require cell-to-cell correspondence. Once cell-level supervision is added ($\lambda = 0.5$ or $1.0$), OT consistently improves performance, showing that matching is mainly needed to provide meaningful per-cell training targets rather than to support bag-level composition learning itself.

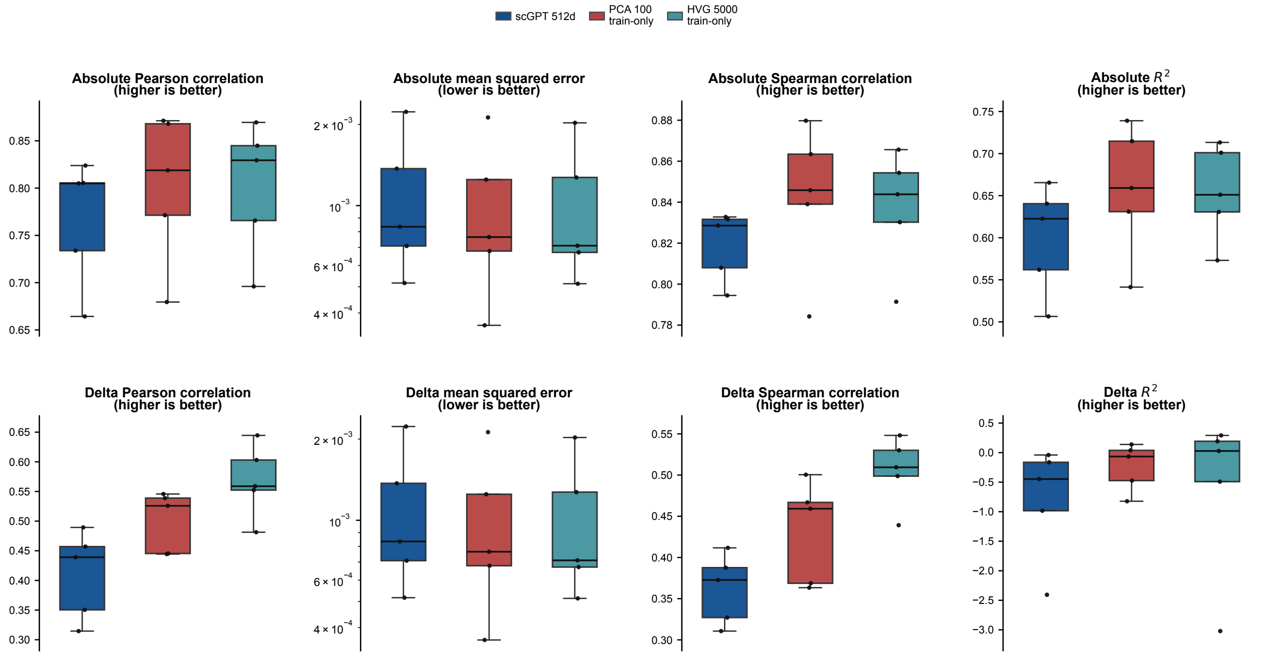


**Fig. S6**: **Cell encoder comparison.** Performance comparison across three cell input representations: top 100 principal components (PCA fitted on the training split), log-normalized counts over the top 5,000 highly variable genes (HVG-5000, training split), and 512-dimensional pretrained scGPT embeddings. Columns report different metrics; the top row shows absolute composition metrics ($\hat{\mathbf{p}}$ vs $\mathbf{p}^*$), the bottom row shows delta-composition metrics ($\Delta\hat{\mathbf{p}}$ vs $\Delta\mathbf{p}^*$). Boxes summarise five-fold cross-validation. The main analysis used pretrained scGPT embeddings as a fixed, transferable cell representation that does not require fitting a dataset-specific feature space. The 100-PC representation matched or exceeded both alternatives across the eight evaluation panels.

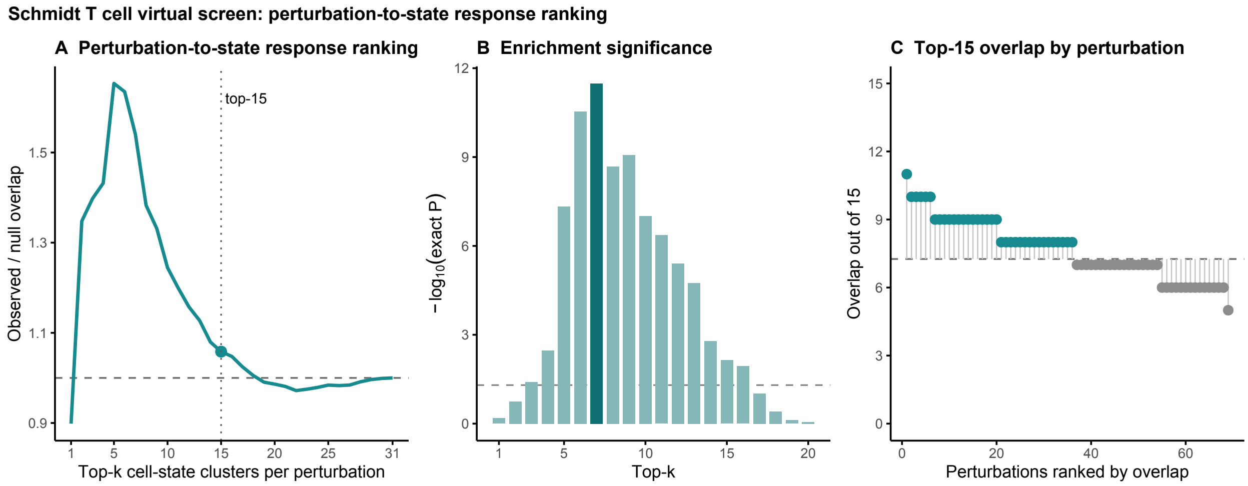


**Fig. S7**: **Perturbation-to-state response ranking in the Schmidt T cell dataset.** For each of 69 evaluated conditions (68 CRISPRa perturbations and the control condition), cell states were ranked by the absolute predicted or observed $\log_2$ fold-change relative to control. At $k = 15$, predicted and observed rankings overlapped in 530 of 1,035 top-ranked slots, compared with 500.8 expected under a stratified hypergeometric null ($P = 0.00725$).

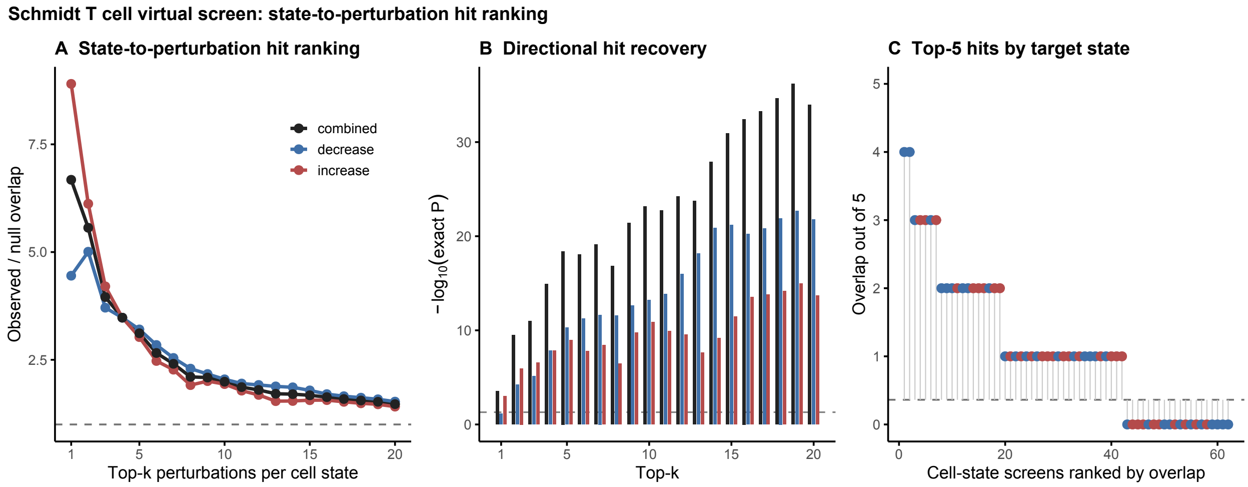


**Fig. S8**: **State-to-perturbation hit ranking in the Schmidt T cell dataset.** For each target cell state and direction of change, 69 candidate conditions (68 CRISPRa perturbations and the control condition) were ranked by predicted and observed signed $\log_2$ fold-change. Predicted top-five conditions recovered 70 of 310 observed top-five hits, compared with 22.5 expected by chance (3.12-fold enrichment; $P = 3.97 \times 10^{-19}$).

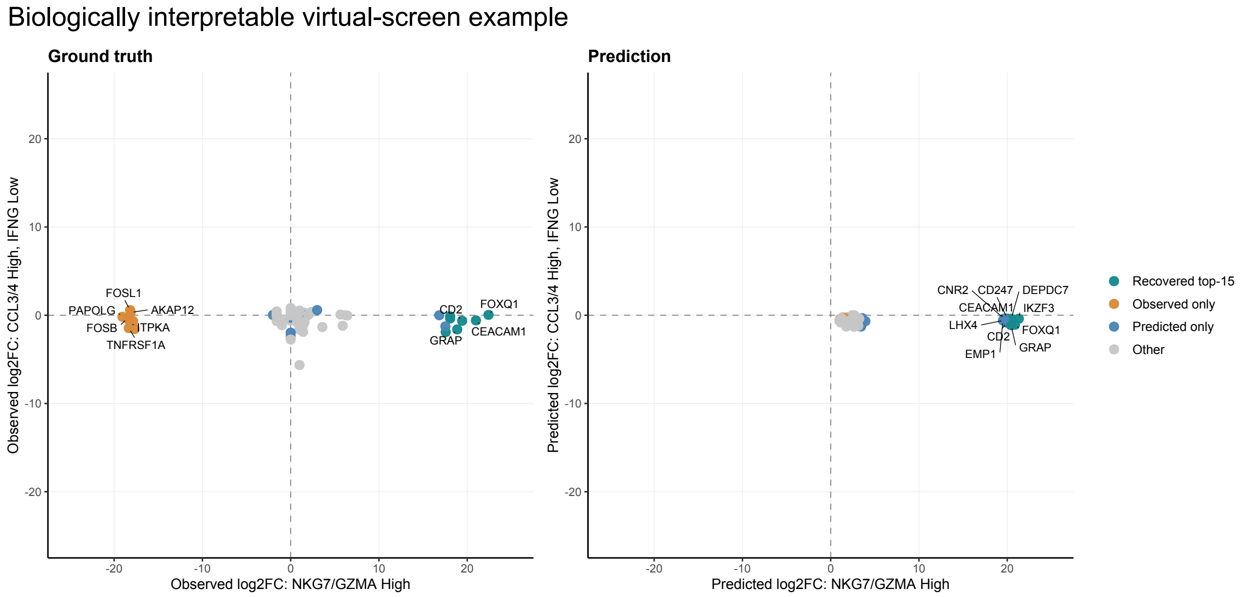


**Fig. S9**: **Illustrative cytotoxic-effector virtual screen.** Example screen along a cytotoxic-effector axis defined by NKG7/GZMA-high and CCL3/4-high, IFNG-low T-cell states. The ranking included 68 CRISPRa perturbations and the control condition. Pop-Corn recovered 7 of 15 observed top conditions in this example, compared with 3.26 expected by chance (2.15-fold enrichment; nominal $P = 0.014$). This example is shown for biological interpretation; the aggregate state-to-perturbation hit ranking in Supplementary Fig. S8 provides the primary statistical evidence.

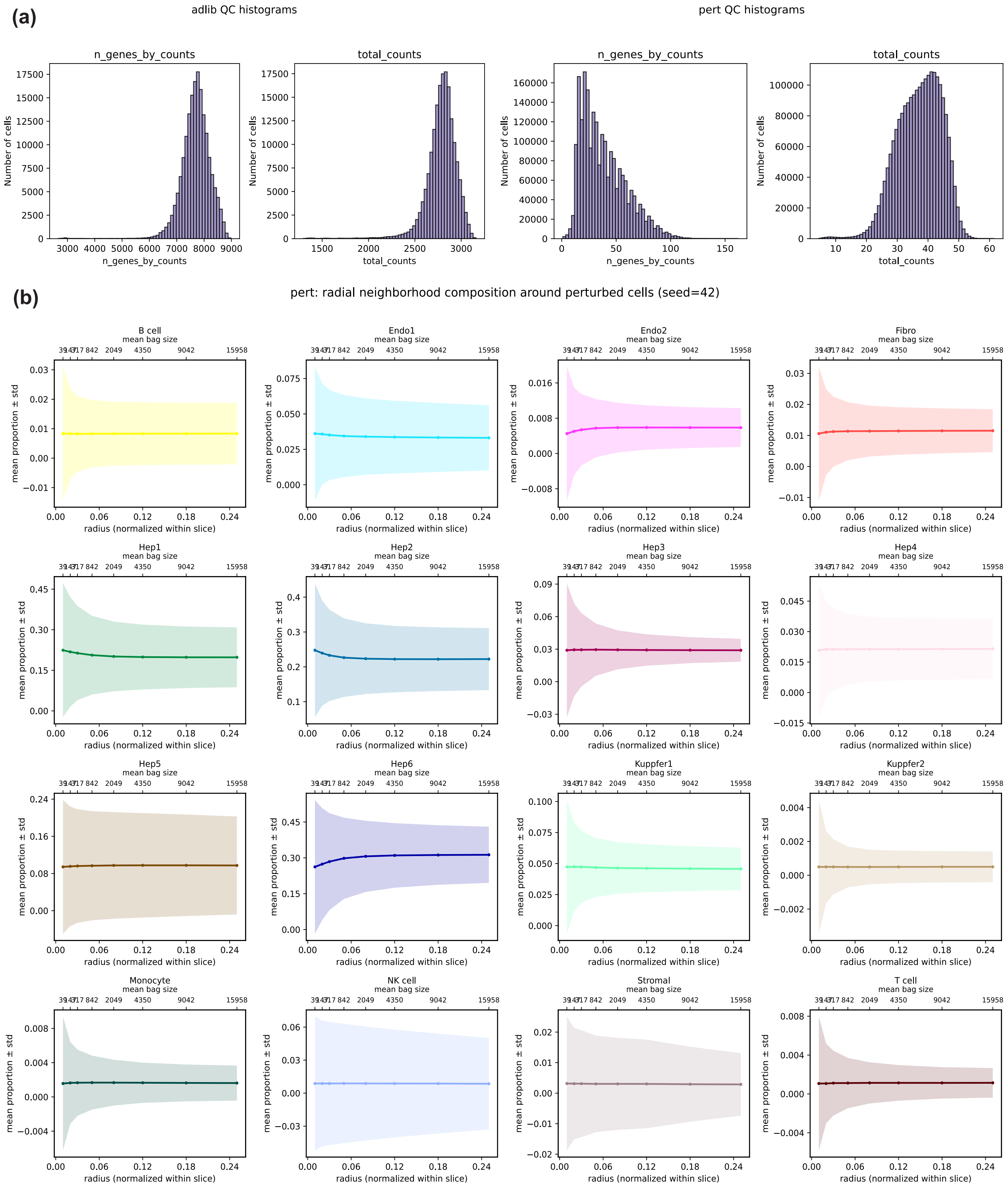


**Fig. S10**: **Spatial Perturb-seq data quality control and bag sampling stability.** Quality control metrics for the spatial CRISPR Perturb-seq dataset (Saunders et al.) and an exploratory analysis of composition stability across radii expressed as fractions of the normalised slice extent. The reported Pop-Corn–Spatial evaluation used 50-$\mu$m niches.

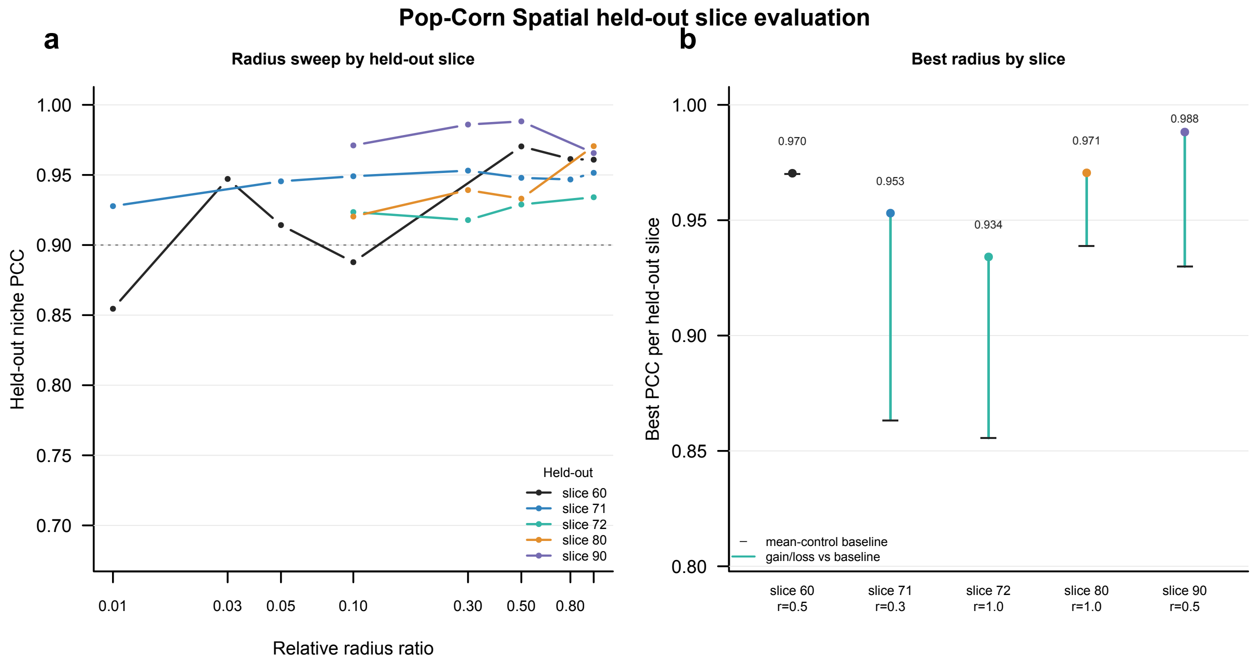


**Fig. S11**: **Spatial bag size comparison.** Pop-Corn Spatial performance across held-out slices and relative spatial bag radii. The radius sweep shows niche-composition PCC for each held-out slice. Best radii varied by held-out slice, with peak PCC values of 0.970, 0.953, 0.934, 0.971 and 0.988 for slices 60, 71, 72, 80 and 90, respectively; the summary panel marks the corresponding mean-control baseline.

### Supplementary Tables

**Table S1**: **Configuration of Pop-Corn for the Schmidt T cell benchmark.** Settings are shared across the five perturbation-held-out cross-validation folds unless otherwise stated.

| Configuration | Setting |
|---|---|
| *Data and evaluation* | |
| Dataset | Primary human T cell CRISPRa Perturb-seq dataset of Schmidt *et al.* |
| Evaluation design | Five-fold cross-validation with perturbation targets held out from training in each fold |
| Random seed | 42 |
| *Input representations and sampling* | |
| Cell representation | 512-dimensional pretrained scGPT cell embeddings |
| Perturbation representation | 2,560-dimensional mean-pooled protein embeddings from ESM-2 model `esm2_t36_3B_UR50D` |
| Training bags | Bootstrap sampling with replacement; sampling fraction 0.3 and bag size $n = \max(1, \lfloor 0.3N \rfloor)$ for a matched pool of $N$ cell pairs |
| Training resamples | 50 bootstrap bags per fold |
| *Model architecture* | |
| Cell encoder | Two-layer feedforward projection to a 64-dimensional shared latent space |
| Perturbation conditioning | Element-wise addition of cell and perturbation representations |
| Set aggregator | One-layer bidirectional transformer with four attention heads, feedforward dimension 128 and dropout 0.1 |
| Composition decoder | Per-cell linear classifier with softmax; bag-level composition obtained by averaging per-cell probabilities |
| *Training and model selection* | |
| Objective | Composition mean-squared error combined with cross-entropy on OT-matched cell labels; cell-level loss weight $\lambda = 0.5$ |
| Optimizer | Adam with learning rate $1 \times 10^{-3}$ |
| Batch size | 32 bags |
| Training budget | Maximum 100 epochs; early-stopping patience of 10 epochs |
| Model selection | Checkpoint with the lowest validation composition mean-squared error |
| *Inference* | |
| Prediction averaging | Mean prediction over $R = 20$ independently sampled control bags per held-out perturbation |

**Table S2: Key notation used in the Pop-Corn formulation.**

| Symbol | Definition |
|---|---|
| $g$ | Genetic perturbation identity |
| $\mathcal{G}_{\text{train}}$ | Perturbations included in the training set |
| $\mathbf{x} \in \mathbb{R}^G$ | Log-normalised expression profile of one cell across $G$ genes |
| $c(x)$ | Annotated cell-type label of cell $x$ |
| $K$ | Number of annotated cell types or states |
| $P_g$ | Distribution of cellular states under perturbation $g$ |
| $P_0$ | Distribution of cellular states in the unperturbed control condition |
| $\pi^g$ | Population cell-type composition under perturbation $g$ |
| $\tilde{\pi}^g$ | Empirical composition estimated from profiled cells under perturbation $g$ |
| $\hat{\pi}^g$ | Cell-type composition predicted by Pop-Corn under perturbation $g$ |
| $\Delta^{K-1}$ | Probability simplex containing non-negative $K$-part compositions that sum to one |
| $n_g$; $n$ | Number of profiled cells under perturbation $g$; number of cells sampled into a bag |
| $\mathcal{B}$ | Finite collection of cells processed jointly by the model, allowing repeated bootstrap draws |
| $F(\mathcal{B}_{\text{ctrl}}, g; \theta)$ | Pop-Corn prediction from a control-cell bag and perturbation identity |
| $\theta$ | Trainable parameters of Pop-Corn |
| $d_h$ | Shared latent dimension of the model |
| $\lambda$ | Weight balancing the bag-level and cell-level training losses |
| $r$ | Radius used to define a niche in a perturbed spatial slice |
| $R$ | Number of independently sampled bags averaged at inference |
